# Pupil- and gaze dynamics track emotion content in naturalistic speech

**DOI:** 10.64898/2026.01.07.698106

**Authors:** Sümeyye Şen Alpay, Christian Keitel, Rosanne H. Timmerman, Anne Keitel

## Abstract

Cortical tracking of speech features is a well-established marker of continuous speech processing, but far less is known about listeners’ ocular responses to speech rhythm. Ocular responses are central to active sensing models in the auditory domain, where motor recruitment guides temporal speech prediction and attention allocation, potentially shaped by non-rhythmic cues, such as emotions. Here, we ask whether listeners’ pupil response and eye movements track the acoustic speech signals and to what extent this tracking is modulated by emotion-related top-down factors, including subjective emotion ratings, mood, and trait empathy. In a validation study (*N* = 100), participants passively listened to two TED talks and intermittently rated segments on valence (negative–positive) and arousal (low–high). This suggested substantial variability in valence and arousal across speech segments in both talks. In the second study (*N* = 41), participants completed the same task while pupillometry and electrooculography (EOG) were recorded. Mutual information was used to quantify speech tracking in pupil dilation, along with horizontal and vertical eye movements. All ocular signals significantly tracked speech at low frequencies. High-arousal speech was associated with stronger pupil tracking but weaker vertical and horizontal EOG tracking. Negative speech valence was linked to stronger tracking in pupil and vertical eye-movement signals. Interactions between the speech-emotion dimensions, as well as their interactions with listeners’ mood, further shaped these effects, giving rise to distinct patterns across ocular measures. Taken together, our findings provide evidence that ocular activity dynamically aligns to the temporal structure of natural speech and that this tracking is sensitive to both stimulus-driven and listener-dependent emotional factors.

## Introduction

Speech is a complex, dynamic acoustic signal that conveys both linguistic and paralinguistic information across time and frequency. Its real-time perception depends on the brain’s ability to parse and interpret these spectro-temporal patterns. Over the past decade, this challenge has driven auditory cognitive neuroscience, with oscillation-based models highlighting speech tracking as a potential mechanism in naturalistic contexts (Atanasova et al., 2025; Doelling et al., 2014; Giraud & Poeppel, 2012). In these models, the quasi-rhythmic temporal structure of speech is highlighted as a core feature in the neural processing of speech. This quasi-rhythmicity in speech is typically quantified by the speech envelope that represents slow amplitude modulations carrying prosodic and syllabic cues (Chi et al., 2005; Rosen, 1992). Electrophysiological studies consistently show that brain activity in the delta (1–4 Hz) and theta (4–8 Hz) bands track the speech envelope (Ahissar & Ahissar, 2005; Doelling et al., 2014; Ghitza, 2017; Gross et al., 2013; Keitel et al., 2018; Luo & Poeppel, 2007; Meyer, 2018; Peelle et al., 2013), putatively facilitating the decoding of acoustic, linguistic, and higher-order information from continuous speech (Ding & Simon, 2014; Obleser & Kayser, 2019). Notably, cortical speech tracking is not purely a bottom-up response. It has been shown to be modulated by top-down cognitive factors such as attention (Lesenfants & Francart, 2020; Rimmele et al., 2015), working-memory load (Hjortkjær et al., 2020), listening effort (Song & Iverson, 2018), and prior expectations (Baltzell et al., 2017). Cortical speech tracking is now widely recognised as an important mechanism for auditory processing, yet the precise neural and computational processes that give rise to it remain to be elucidated.

Tracking of speech rhythm appears to extend beyond brain rhythms, manifesting also in other bodily rhythms. Recent work shows that eye movements, blinks, and pupil dynamics align with speech features in synthesised speech (Jin et al., 2018), short sentences (Gehmacher et al., 2024), and naturalistic continuous speech (Holtze et al., 2023; Madsen & Parra, 2024; Schubert et al., 2025), with a consistent link between this tracking and attentional mechanisms. Specifically, pupil response stands out as a temporally rich signal in recent oscillation-based frameworks (Keitel, et al., 2025; Liu et al., 2025), governed by autonomic nervous system activity, with sympathetic and parasympathetic influences driving pupil dilation and constriction respectively (Mathôt, 2018). Ongoing fluctuations in pupil dilation reflect arousal, stemming from subcortical neuromodulatory systems (Costa & Rudebeck, 2016), primarily the locus coeruleus-norepinephrine pathway, which is often characterised by phasic (i.e. event-related response) and tonic (i.e. task-free, spontaneous fluctuations) modes (Aston-Jones & Cohen, 2005; Liu et al., 2025). On the other hand, eye-movement based speech tracking has been linked to cortical tracking of speech, implying motor and attentional engagement in auditory processing (Gehmacher et al., 2024; Schubert et al., 2025). Together, these findings suggest that eye responses support auditory speech perception and may serve as a proxy for sensorimotor engagement. Still, further research is needed to establish the reliability and generalisability of these measures and to examine top-down influences beyond attention, as well as their potential interactions, in ocular speech tracking.

One such influence may be the emotional content of speech, which is particularly relevant as emotions are inherent at nearly every level of spoken communication, from phonetic detail to semantic, grammatical, and discourse cues (Majid, 2012). Research on emotional speech processing has mostly focused on prosody, that is, the suprasegmental modulation of tone, stress, and rhythm, which serves both linguistic and emotional functions (Myers et al., 2019; Paulmann, 2016). On the other hand, speech tracking research has largely focused on the linguistic functions of prosody, neglecting its social and emotional dimensions (see: Rimmele et al., 2021; Teoh et al., 2019). Yet linguistic and emotional functions of prosody are closely intertwined with shared acoustic parameters and neural pathways, (Paulmann, 2016; Pell & Kotz, 2021; Themistocleous, 2025), making consideration of the emotional aspects equally essential. Indeed, it has already been suggested that connecting patterns of envelope tracking to the perception of emotions could offer practical insights for clinical populations characterised by prosodic difficulties, as well as for typically developing individuals (Myers et al., 2019). Here, we empirically test the relationship between the emotion content in continuous speech and ocular responses.

Emotions are known to have preferential access to attentional resources, thereby facilitating adaptive behaviours in complex environments (Ceravolo et al., 2016; Guex et al., 2022; Ho et al., 2015; Pessoa, 2008; Vuilleumier, 2005). Behavioural studies consistently report enhanced performance in a variety of tasks, such as faster reaction times and greater accuracy when processing emotional speech compared to neutral speech (Gordon & Ancheta, 2017; Kim & Sumner, 2017; Lu et al., 2021; Olano et al., 2020; Wurm et al., 2001). Neuroimaging work further reveals distinct electrophysiological and hemodynamic patterns for emotional prosody (Belyk & Brown, 2014; Grandjean et al., 2005; Mauchand & Zhang, 2023; Paulmann et al., 2011; Schirmer & Kotz, 2006), suggesting emotion may impact early auditory processing. Taken together, these findings make emotions a strong candidate for modulating speech tracking at both neural and physiological levels.

The present study was guided by two complementary aims. First, we sought to extend existing evidence for speech rhythm tracking by ocular responses during passive listening to continuous speech, focusing specifically on pupil dilation and eye movements. Second, we tested whether and how emotional factors relate to ocular speech tracking. To address these aims, we conducted two studies: an initial validation study that characterised the emotional arousal and valence of our naturalistic speech stimuli in a large sample (*N* = 100), followed by the main experiment (preregistration: https://doi.org/10.17605/OSF.IO/X2SRW), in which participants (*N* = 41) listened to the same material while we recorded electrooculography and pupillometry. We then quantified the coupling between the speech signal and each ocular measure and examined its relationship to subjective ratings of emotional arousal and valence, listeners’ mood and empathy. We tested the hypothesis that higher emotional arousal would enhance ocular speech tracking, potentially through increased attention. We did not make a priori hypothesis about the effects of valence and its potential interactions with arousal since less is known about how these factors shape speech processing in naturalistic settings.

### Study 1: Validating emotional speech stimuli

In Study 1, we characterised the emotional and acoustic features of our naturalistic speech excerpts. We chose two TED talks as stimuli (Talk 1: 13 min; Talk 2: 12 min) that varied in their emotional content during a single talk. Participants were asked to intermittently rate the stimuli on arousal and valence scales. Note that several validated emotional speech databases are currently available in the literature. However, these were not suitable for our study’s aims due to a range of limitations such as speaker accent, stimulus length, reduced naturalness of speech delivery, and copyright restrictions.

Our analyses addressed five research questions. First, we examined whether emotional responses differed between the two TED Talks, to assess the influence of content and speaker-specific delivery on listeners’ emotional evaluations. Second, we assessed the consistency of participants’ ratings to evaluate the reliability of the emotional responses elicited by the speech. Third, we analysed whether specific acoustic features were associated with participants’ arousal and valence ratings. Fourth, we examined the relationship between arousal and valence to better understand how these affective dimensions co-occur in natural speech. Finally, we investigated whether individual differences, including self-reported mood and trait empathy, were related to variation in emotional ratings.

The treatment of participants adhered to the principles of the Declaration of Helsinki. Written informed consent was obtained from all participants. The study was approved by the School of Social Sciences Ethics Committee at the University of Dundee (Approval No: UoD-SHSL-PSY-PG-2023-164). Participants received compensation in the form of monetary payment (£5) or course credit, depending on their preference.

## Materials and methods

### Participants

A sample of *N* = 100 volunteers (77 female, 20 male, and 3 non-binary participants; *M*_age_= 24.5 years, age range = 17–48 years, *SD_age_*= 6.84) were recruited via the department’s participant pool and advertisements. All participants reported advanced English proficiency and no prior diagnoses of speech or language disorders, neurodevelopmental conditions, or psychological or neurological disorders. Participants completed the *Quick Hearing Test* (QHT; (Koike et al., 1994) to assess their self-reported hearing ability. Four participants obtained scores (28,28,32,38 out of 60) that may suggest slightly diminished hearing (hearing test is recommended for scores ≥ 20). Participants completed additional self-report measures, including the *Brief Mood Introspection Scale* (BMIS; (Mayer & Gaschke, 1988), which assesses current mood, and the *Toronto Empathy Questionnaire* (TEQ; (Spreng et al., 2009), measuring emotional understanding and responsiveness to others. Participants’ average mood score was *M* = 49.2 (*SD* = 5.78) on the valence dimension and *M* = 27.8 (*SD* = 3.75) on the arousal dimension, while the average total empathy score was *M* = 50.8 (*SD* = 6.50).

### Speech stimuli

We used two audio recordings of TED Talks delivered by female speakers with British accents (Talk 1: “You can be fat and happy” by Sofie Hagen; Talk 2: “Breaking out of concrete: six ways out of depression” by Bethany Rose). We edited the original recordings to shorten long silences, audience reactions, such as laughter and applause, and segmented each talk into speech chunks ranging from 7.27 s to 22.35 s (*M* = 13.90, *SD* = 3.81 s). We ensured that speech chunks were emotionally coherent. This process resulted in 101 distinct speech chunks (N_talk1_ = 55, N_talk2_ = 46), each containing between one and six sentences of varying structure.

We presented all speech stimuli using PsychToolbox 3.0.17 (Kleiner et al., 2007), delivering audio playback with high-quality wired headphones (Sennheiser HD-25, 70Ω). Additionally, we amplitude-normalised the stimuli to maintain consistent volume levels across all excerpts. Audio files had a sampling rate of 44,100 Hz. The speech stimuli are publicly available on the OSF server (https://osf.io/dsvzh/files/osfstorage).

### Procedure and task

Participants were tested in a quiet room while seated at an approximate distance of 65 cm from the screen. They were instructed to fixate on the circle presented at the centre of the screen, to listen attentively to the speech stimuli, and to rate each chunk on arousal and valence scales. To familiarise themselves with the task, they first completed a practice trial in which they listened to three speech chunks from a TED Talk that was not included in the main task. They were allowed to adjust the volume to their preferred level before the main task.

The main task consisted of two blocks, each corresponding to one of the TED Talks. The order of the blocks was randomised, whereas the order of the speech segments was kept constant to preserve the natural flow of the narrative. A self-paced break was provided between the two blocks. Participants rated their own felt emotions on visual analogue scales for valence and arousal separately (see *Figure 1*). For valence ratings, participants answered the question “How did the last speech chunk make you feel?” using a visual analogue scale from −100 to 100, anchored by “Very negative” and “Very positive”. Arousal was measured with the question “How intense was your emotional response to the speech chunk?”, accompanied by a visual analogue scale ranging from 0 to 100 with anchors “Low arousal” and “High arousal”. Participants indicated their response by placing a vertical line on the scale through a mouse click. At the end of each talk, participants completed a familiarity rating to assess whether they had encountered the speech material before. The session, including instructions, questionnaires, and two passive listening blocks, lasted approximately one hour.

**Figure 1.**
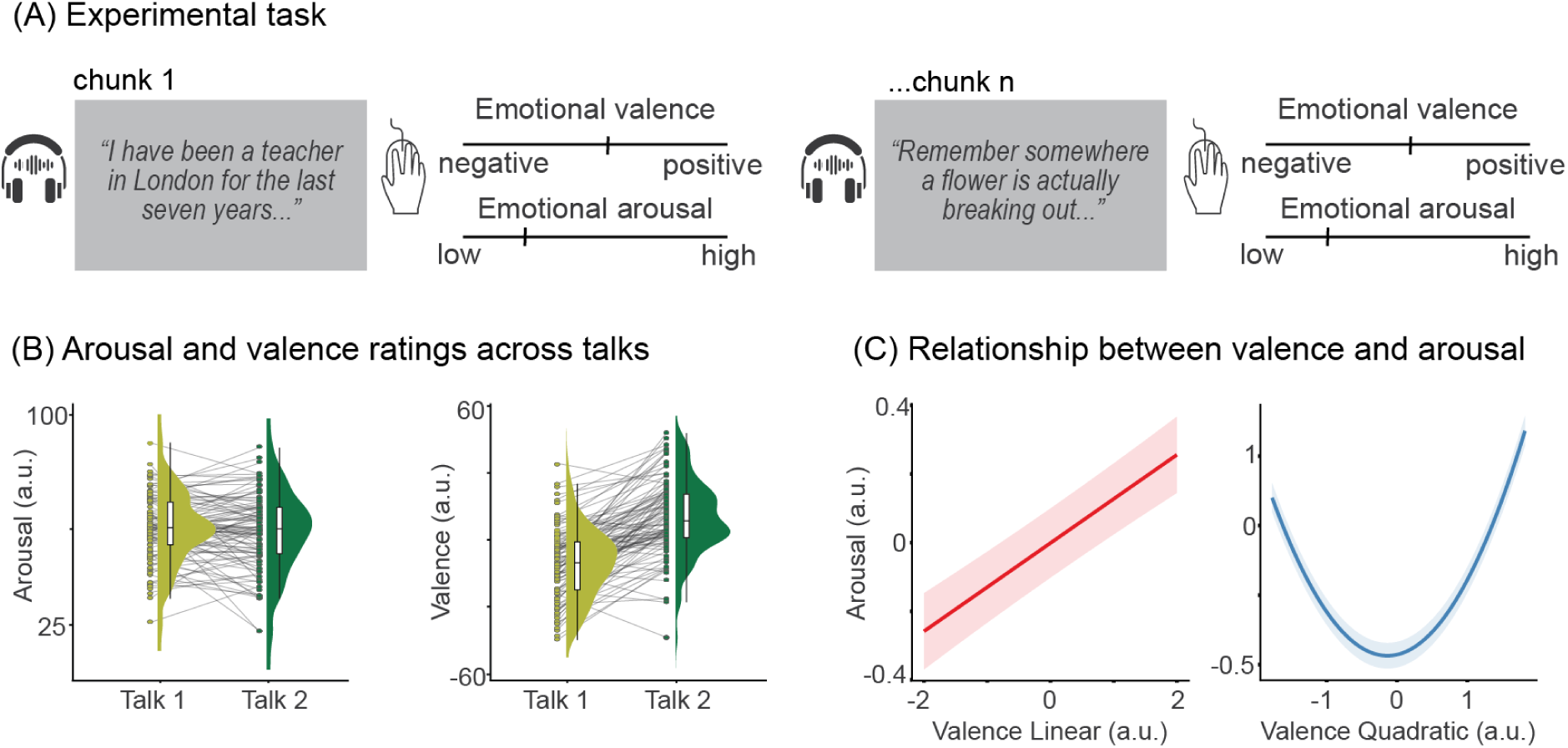
Overview of the experimental task and measures. (A) A passive listening and rating task was used in Study 1. Participant listened to speech chunks (between 7 and 22 s) while fixating on a circle with the same luminance as the background. Then they rated each chunk on arousal and valence ratings. (B) Raincloud plots showing mean arousal and valence ratings across speech chunks for Talk 1 and Talk 2 (*N* = 100). Each point represents an individual participant’s mean rating; half-violin plots illustrate the kernel density of these means. Light grey lines connect each participant’s mean emotion ratings between talks, emphasising within-subject changes. (C) The linear and quadratic relationship between emotion ratings. The figure was derived from a linear mixed-effects model with arousal as outcome and displays the predicted arousal values from the model (y-axis) as a function of the valence linear (left) and valence quadratic (right) (x-axis), with shaded areas representing 95% confidence intervals. The model revealed significant main effects of valence (both linear and quadratic) on arousal ratings.

### Emotion model

Although there is not yet consensus in the literature on how emotions should be modelled (Hamann, 2012), we here adopted a well-established dimensional approach to conceptualise emotions. In the dimensional approach, an emotion can be represented by a specific point within a continuum defined by a small set of underlying dimensions, which may be more appropriate for a more nuanced assessment of emotional experience compared to discrete models (Cowie & Cornelius, 2003; Laukka et al., 2005). Specifically, we used the circumplex model of affect, which postulates that emotions can be mapped into two orthogonal dimensions: valence, referring to the degree of pleasantness of an emotion, and arousal, reflecting its level of intensity or activation (Russell, 1980).

### Acoustic feature extraction

We quantified the acoustic characteristics of each speech chunk by extracting features with the openSMILE toolkit (Eyben et al., 2010) in Python. We applied the extended Geneva Minimalistic Acoustic Parameter Set (eGeMAPS; (Eyben et al., 2015), which includes standardised parameters commonly used for affective speech processing. This extraction yielded 88 acoustic features per speech chunk, belonging to either frequency, energy, spectral, or temporal domains.

To reduce dimensionality and address multicollinearity among predictors, we performed a Principal Component Analysis (PCA) with varimax rotation on the full set of extracted features (Jolliffe & Cadima, 2016). We retained the first two principal components, which explained 39% and 11% of the variance, respectively, resulting in a cumulative explained variance of approximately 50%. We then selected the five acoustic features with the highest absolute loadings from each component for use as predictors in subsequent models (see Kamiloğlu et al., 2020 for a similar approach). These ten features reflected both core prosodic attributes (e.g., pitch variability, loudness, jitter) and spectral shape characteristics (e.g., Mel-Frequency Cepstral Coefficients (MFCC), spectral slope). Table 1 provides a list of the selected features and their factor loadings.

**Table 1.** Acoustic features with highest loadings on the first two components. F1, F2. F3: the ratio of the energy of the spectral harmonic peak at the first, second, third formant’s centre frequency to the energy of the spectral peak at F0. MFCC4 (M): Arithmetic mean of the fourth MFCC across all frames, reflecting midlevel spectral contour. Loudness (M): Arithmetic mean of perceived intensity values computed from an auditory weighted Melband spectrum. F₀ (SD): Standard deviation of the fundamental frequency across voiced frames, indicating pitch variability. Harmonics-to-Noise Ratio (M): Arithmetic mean of the ratio between harmonic energy and noise energy, indicating voice periodicity. Alpha Ratio (SD): Standard deviation of the ratio between summed spectral energy, capturing variability in low vs high frequency balance. Jitter (M): Arithmetic mean of cycle-to-cycle period length deviations, reflecting glottal vibration irregularity. Spectral Slope (M): Arithmetic mean of the linear regression slope of the log power spectrum. Computational details and the relevance of each parameter are provided in Eyben et al. (2015).

|  | Acoustic feature | Group | Factor loadings |
| --- | --- | --- | --- |
| Component 1 | F<sub>3</sub> relative energy (M) | Spectral | .95 |
|  | F<sub>2</sub> relative energy (M) | Spectral | .95 |
|  | F<sub>1</sub> relative energy (M) | Spectral | .91 |
|  | MFCC4 (M) | Spectral | .85 |
|  | Loudness (M) | Temporal | .84 |
| Component 2 | F<sub>0</sub> (SD) | Frequency | .85 |
|  | Harmonics-to-noise ratio (M) | Energy | .74 |
|  | Alpha ratio (SD) | Spectral | .71 |
|  | Jitter (M) | Frequency | .69 |
|  | Spectral slope (M) | Spectral | .69 |

### Statistical analyses

Unless otherwise noted, subsequent relationships were tested using Linear Mixed-Effects Models (LME) in R (v4.4.2; *R Core Team*, 2024) using the *lme4* (Bates et al., 2015) package. The significance of fixed effects was evaluated using the *lmerTest* package (Kuznetsova et al., 2017), which applies Satterthwaite’s approximation to estimate degrees of freedom. Model visualisations were generated with the *sjPlot* package (Lüdecke et al., 2025). To control for multiple comparisons, *p* values were adjusted using the false discovery rate (FDR) (Benjamini & Hochberg, 1995). The significance of main and interaction effects was further assessed using likelihood-ratio tests comparing nested models with and without the fixed effects of interest (Gelman & Hill, 2007). Model parameters were estimated using an unrestricted maximum likelihood optimisation procedure. Continuous predictors were standardised (z-scored). Assumptions for multilevel analyses (linearity, homogeneity of variance, multicollinearity, and normality of residuals) were visually inspected via diagnostic plots. A significance level of α = 0.05 was set for all the models.

## Results

### Relationship between emotional arousal and valence

To explore the relationship between arousal and valence ratings, we fitted a model with arousal ratings as the outcome and valence as a fixed effect, including both linear and quadratic terms to capture potential non-linear associations. We found significant main effects of both the linear term (*β* = 11.33, 95% CI [9.30, 13.36], *p* < .001, *p*_FDR_ < .001) and the quadratic term (β = 49.19, 95% CI [47.04, 51.34], *p* < .001, *p*_FDR_ < .001). The linear valence model fit significantly better than the null model (χ²(1) = 114.1, *p* < .001) and the quadratic model provided an additional significant improvement over the linear model, χ²(1) = 1773.8, *p* < .001. The linear relationship indicates that more positive valence was associated with higher arousal, while the quadratic effect suggests that both highly positive and highly negative speech chunks elicited higher arousal compared to neutral ones. The quadratic association, illustrated with a regression curve, can be seen in the Supplementary Materials (Figure S2).

### Relationship between emotional responses to speech and acoustic features

To examine whether acoustic features predicted subjective ratings of emotions, we fitted two separate linear mixed-effects models for arousal and valence. Both models included the ten PCA-selected acoustic predictors as fixed effects, and arousal and valence as outcome variables. Since the formant frequency measures showed severe collinearity (VIF > 30), the F_2_ and F_3_ mean variables were removed from the models. After this step, multicollinearity among the remaining predictors fell within acceptable limits (VIF < 6).

For valence, only loudness significantly predicted participants’ ratings (*β* = 0.32, 95% CI [0.03, 0.62], *p* = .033). This effect did not remain significant after controlling for multiple comparisons (*p*_FDR_ = .224); however, the full model provided a significantly better fit than the null model (χ²(8) = 10.21, *p* = .014). Loudness reflects the intensity of the speech signal, which is closely related to the amplitude of vocal output. The positive coefficient indicates that a louder voice tended to be associated with more positive speech.

For arousal, the main effect of F₀(SD) significantly predicted ratings (*β* = −0.09, 95% CI [−0.16, −0.02], *p* = .016), as well as the main effect of F_1_ mean (*β* = −0.11, 95% CI [−0.20, −0.02], *p* = .015). However, neither effect survived after FDR correction (both *p*_FDR_ = .082). Notably, the arousal model did not show a significant improvement in fit over the null model containing only random effects (χ²(12) = 12.75, *p* = .12). F₀(SD) reflects pitch variability, and the negative association suggests that greater variability was related to lower perceived arousal, although the effect does not survive correction for multiple comparisons. Similarly, F_1_ mean, a parameter perceived as vowel sound, was negatively associated with speech arousal, but this effect also did not remain significant.

### Effects of individual differences on emotion ratings

To examine whether participants’ subjective arousal and valence ratings were influenced by individual differences, two separate linear mixed-effects models were fitted with empathy and mood as fixed effects.

For valence, participants who were in a more positive mood (valence dimension) tended to rate the speech material more positive (*β* = 0.06, 95% CI [0.01, 0.11], *p* = .011, *p*_FDR_ = .051). Mood arousal (*p* = .14, *p*_FDR_ = .28) and empathy (*p* = .83, *p*_FDR_ = 1) were not significantly related to valence ratings. The overall model did not show an improved fit when compared to the null model (*χ²*(3) = 7.74, *p* = .052).

For arousal, higher empathy scores predicted increased ratings (*β* = 0.12, 95% CI [0.04, 0.21], *p* = .005, *p*_FDR_ = .023). In contrast, neither mood valence (*p* = .56, *p*_FDR_ = 1) nor mood arousal (p = .87, *p*_FDR_ = 1) showed significant effects. Model comparison indicated that including empathy and mood predictors significantly improved model fit relative to the null model (*χ²*(3) = 8.76, *p* = .033).

### Interrater reliability

To quantify the reliability of emotion ratings across participants, we calculated intraclass correlation coefficients (ICCs) separately for arousal and valence. Specifically, we used the ICC(2,k) model, a two-way random-effects model that estimates the consistency of average ratings across multiple raters sampled from a larger population (Koo and Li, 2016). This model is appropriate when the same set of raters evaluate all items, and the goal is to generalise to other raters. We interpreted ICC values according to the thresholds proposed by Koo and Li (2016): poor (< .50), moderate (.50–.75), good (.75–.90), and excellent (> .90). Valence ratings demonstrated excellent interrater reliability, ICC(2,k) = .99, 95% CI [0.99, 0.99], *F*(100, 9900) = 98, *p* < .001. Arousal ratings showed good reliability, ICC(2,k) = .87, 95% CI [0.83, 0.90], *F*(100, 9900) = 9.4, *p* < .001.

### Differences in emotion ratings between talks

We tested whether emotional responses differed across the two TED talks, using separate models for arousal and valence ratings as outcome variables and *Talk* (Talk 1 vs. Talk 2) as a fixed effect. The model including *Talk* significantly improved model fit for valence ratings, *χ²*(1) = 7.42, *p* = .006, with more positive valence ratings for Talk 2 (*β* = 0.37, 95% CI [0.10, 0.63], *p* = .006, *p*_FDR_ = .014). In contrast, the main effect of *Talk* on arousal was not significant (*p* = .39, *p*_FDR_ = .715). The chunk-level distributions of arousal and valence ratings are provided in Supplementary Materials (Figure S1).

## Discussion

In Study 1, we demonstrated that our stimulus set can reliably elicit differentiated emotional responses, evidenced by strong interrater agreement for valence and good interrater agreement for arousal ratings. This supports the robustness of our rating procedure and justifies the use of participant ratings in the subsequent study. Arousal ratings did not differ across the two talks, whereas valence ratings were higher for Talk 2, suggesting a difference in emotional tone that may reflect variations in content or delivery style.

We also observed both linear and quadratic relationships between valence and arousal ratings (see Supplementary Materials, Figure S2). The linear trend suggests that more positive speech chunks were generally perceived as more arousing. This is consistent with the proposal that these two dimensions are independent, where positive emotions like excitement are high in arousal, and negative ones like sadness are low (Kuppens et al., 2013; Russell, 1980). However, the significant quadratic effect suggests that a U-shaped relation where both positive and negative extremes elicit stronger arousal than neutral speech. This pattern has been documented in behavioural studies (Kanske & Kotz, 2010), and are supported by neuroimaging findings, which identify partially distinct neural responses to valence and arousal within the voice perception network (Bestelmeyer et al., 2017).

Despite substantial efforts to identify acoustic markers of emotion (Bachorowski, 1999; Banse & Scherer, 1996; Cowie & Cornelius, 2003; Goudbeek & Scherer, 2010; Laukka et al., 2005; Patel et al., 2011; Schröder, 2001), the precise mapping between vocal features and emotional states remains inconsistent and, at times, contradictory (Larrouy-Maestri et al., 2025). While high-arousal emotions are typically associated with higher pitch and greater pitch variability in speech (Patel et al., 2011), our results revealed the opposite pattern: greater pitch variability was linked to lower perceived arousal. One possible interpretation is that exaggerated pitch fluctuations in naturalistic speech may have been perceived as less intense or even less authentic, thereby reducing arousal judgments.

Our results suggest that loudness was positively associated with valence ratings, indicating that louder voices were evaluated as more emotionally positive. Loudness refers to the subjective perception of sound intensity, and individuals can perceive the same sound as louder or softer depending on several factors. The evidence linking emotional valence to the loudness of auditory stimuli is mixed (Bigras et al., 2024; Weninger et al., 2013; Yang et al., 2018). Within the speech domain, loudness is primarily considered an indicator of high arousal (Juslin & Laukka, 2003), and empirical evidence for its relationship with valence is relatively limited. One available study that analysed multiple datasets reported a negative association between loudness and valence (Weninger et al., 2013). Our results contrast with this finding by showing a weak positive association. This is likely due to the specific speech stimuli used in the present study, where positive expressions coincided with a raised voice. Taken together, our present findings point to pitch (F₀) variability, first formant (F_1_), and intensity as potential indicators of emotional arousal and valence in the here used continuous, natural speech stimuli.

Individual differences further shaped emotional responses. Participants with higher empathy reported stronger arousal to emotional speech, in line with prior findings suggesting that empathy enhances sensitivity to emotional cues (Maffei et al., 2019; Martínez-Velázquez et al., 2020; Nebi et al., 2022; Neves et al., 2018; Wallmark et al., 2018). Our findings extend this literature by showing that empathy also shapes how listeners perceive emotional arousal in continuous speech. One explanation is that empathic listeners may be more susceptible to emotional contagiousness, resonating more deeply with speakers’ emotional arousal. While such a mechanism might also be expected to influence valence perception, the relatively narrower range of valence ratings (see Fig 1. C–D) likely constrained the detection of individual differences, limiting the observable impact of empathy on this dimension.

The mood congruence effect posits that individuals’ emotional state can bias the perception and evaluation of information (Bower, 1981). In support of this, we found that valence evaluation of speech was predicted by participants’ mood; positive mood was associated with more positive ratings, whereas negative mood was associated with more negative ratings, complementing existing evidence in the auditory domain (Egidi & Nusbaum, 2012; Hofbauer & Rodriguez, 2023; Houston & Haddock, 2007; Nygaard & Queen, 2008).

Overall, these validated speech stimuli provide us a solid foundation for Study 2, which investigated the effect of emotions on ocular speech tracking.

### Study 2: Ocular responses to speech rhythm: Investigating the effects of perceived emotions

In Study 2, we aimed to examine how ocular physiological signals (pupil dilation, horizontal and vertical EOG) interact with the quasi-rhythmic structure of continuous speech and how emotions shape this tracking. Recent findings indicate that, in addition to the well-established alignment between speech acoustics and cortical rhythms, ocular dynamics may also track speech (Gehmacher et al., 2024; Holtze et al., 2023; Jin et al., 2018; Madsen & Parra, 2024; Schubert et al., 2025). However, available evidence is currently insufficient and inconsistent to determine whether speech tracking generalises to different ocular signals. Moreover, it is unlikely that such tracking operates independently of stimulus characteristics, listening environments, as well as cognitive and affective influences. While attention has been shown to modulate the temporal alignment between speech and pupil dilation (Jin et al., 2018), as well as eye movements (Gehmacher et al., 2024; Schubert et al., 2025), it is unclear how other factors may influence ocular speech tracking. Here, we investigated how the perceived emotional characteristics of the speech material and other emotion-related factors (listener’s own mood state and trait empathy) were linked to ocular speech tracking.

There are (at least) four reasons to expect that emotions may shape speech tracking in ocular responses. First, emotional expressive speech shows more pronounced and frequent acoustic variability than emotionally neutral speech (Banse & Scherer, 1996; Mullennix et al., 2002). Such emotion-induced changes in the speech signal are likely to influence envelope tracking. Second, there is robust evidence that emotionally expressive speech in different forms (isolated words or short sentences) is processed differently from neutral speech at both neural and behavioural levels (Gordon & Ancheta, 2017; Pell & Kotz, 2021; Pinheiro et al., 2017). Third, the ocular measures we used have been associated with auditory emotional processes. Specifically, emotionally arousing stimuli have been shown to elicit greater pupillary response, potentially through increased autonomic nervous system activity (Bradley et al., 2008; Gingras et al., 2015; Jürgens et al., 2018; Kirwan et al., 2025; Partala & Surakka, 2003). Beyond basic autonomic responses to arousing stimuli, some evidence suggests that pupil dilation is sensitive to emotional valence (Jürgens et al., 2018; Kuchinke et al., 2011; Merchie et al., 2024) and emotional authenticity (Cosme et al., 2021), and is further associated with audiovisual integration (Arias Sarah et al., 2023) as well as emotion-related decision processes (Oliva & Anikin, 2018) in response to vocal emotional stimuli (see Zekveld et al., 2018 for an overview of pupillometry studies in auditory domain). Direct evidence for the role of eye movements in the vocal emotion literature is scarce, with most support coming from cross-modal studies showing that emotional prosody can facilitate gaze behaviour in visual tasks (Gerdes et al., 2021; Paulmann et al., 2012; Rigoulot & Pell, 2012). Finally, emotional stimuli are speculated to be prioritised in attention (Pessoa, 2008), implying that emotion-related changes in attentional mechanisms could shape the extent to which ocular signals align with the speech signal.

The main analyses were preregistered prior to data analysis (https://doi.org/10.17605/OSF.IO/X2SRW). However, there were some small deviations from the preregistration as we encountered new analytic developments, which are described below where relevant.

## Materials and methods

### Participants

A total of N = 44 participants completed the experiment. Data from three participants were excluded for the following reasons: one withdrew from the study, one reported intentionally providing neutral ratings for every chunk, and for one, technical issues resulted in a failure to save electrooculographic data. The final sample consisted of 41 participants (28 female, 13 male; *M*_age_ = 23.2 years, age range = 17–39 years, *SD_age_*= 5.56; 36 right-handed, 3 left-handed, 2 cross-dominant). Thirty-nine participants provided complete datasets; for the remaining two, only partial data were analysed: one contributed only EOG data due to missing pupil data, and the other completed only one talk.

All participants were native English speakers. They reported no prior diagnosis of speech or language disorders, neurodevelopmental conditions, or psychological or neurological disorders. Self-report measures were identical to Study 1, including the *Quick Hearing Test* ( *QHT*; Koike et al., 1994), *Brief Mood Introspection Scale* ( *BMIS*; Mayer & Gaschke, 1988), and *Toronto Empathy Questionnaire* (*TEQ*; Spreng et al., 2009). 39 participants had normal hearing, while two obtained scores (23 and 24 out of 60) that may suggest slightly diminished hearing (hearing test is recommended for scores ≥ 20). Participants’ average mood score on the valence dimension was *M* = 48.8 (*SD* = 5.71), the average mood score on the arousal dimension was *M* = 26.4 (*SD* = 3.69), and average empathy score was *M* = 50.2 (*SD* = 5.85).

All aspects of the study adhered to the principles of the Declaration of Helsinki. Written informed consent was obtained from all participants. The study was approved by the School of Social Sciences Ethics Committee at the University of Dundee (Approval No: UoD-SHSL-PSY-PG-2023-164). Participants received compensation in the form of monetary payment (£15) or course credit, depending on their preference.

### Stimuli and task

We used the same two TED talks as speech stimuli as in the first study. Speech rate was computed in Praat (Version 6.0.18; Boerma & Weenink, 2007), with 3.50 Hz for the first talk and 3.22 Hz for the second talk (de Jong & Wempe, 2009). The procedure and task were also identical to that in Study 1 (*Figure 1A*). However, in Study 2, we recorded eye movements and pupil size throughout the experiment using electrooculography (EOG) and pupillometry, respectively (*Figure 2A*). The session, including setup, instructions, questionnaires, and two passive listening blocks, lasted approximately 1.5 hours.

**Figure 2.**
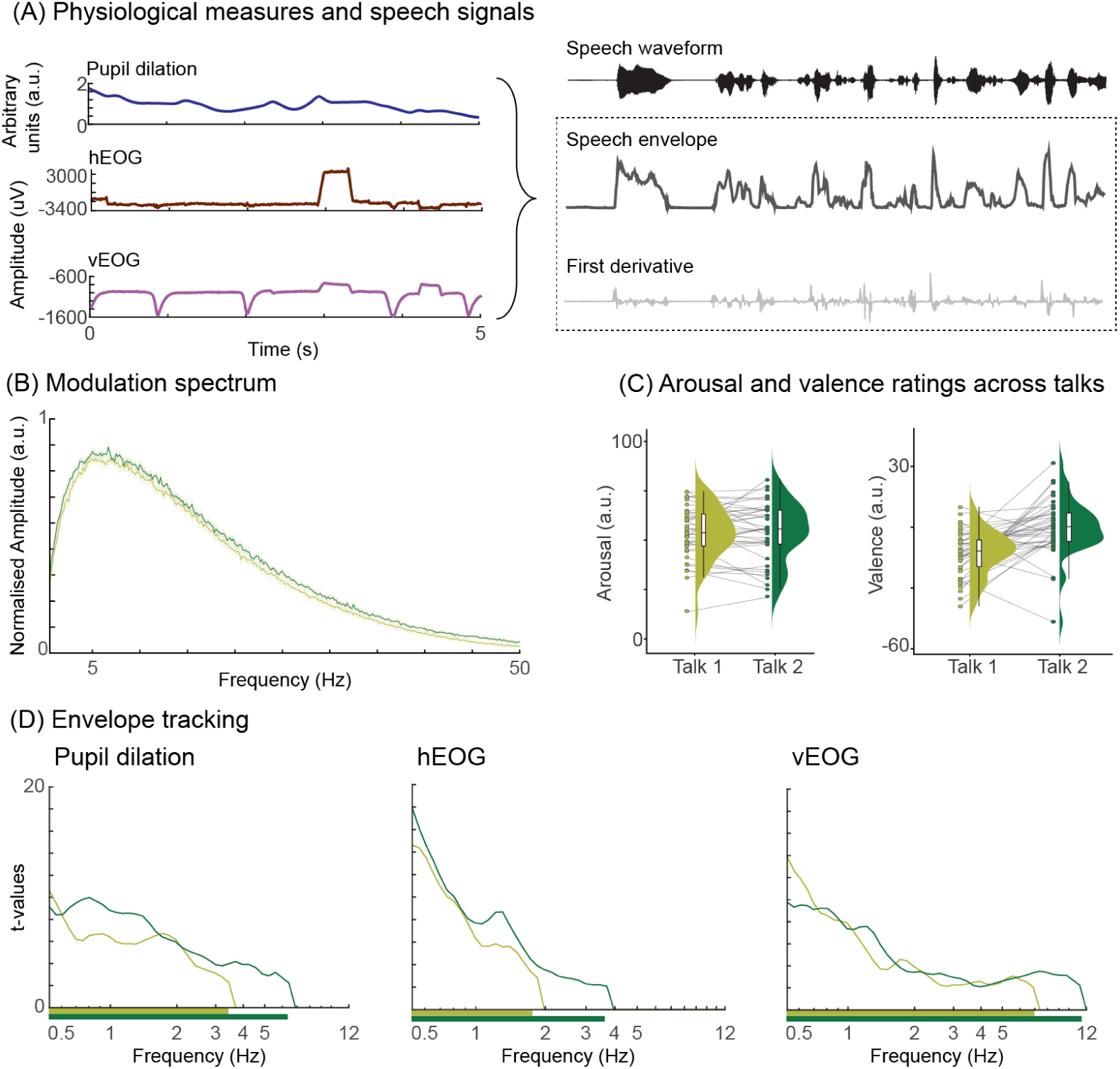
Overview of the experimental task and measures. (A) In Study 2, participants completed the same passive listening and rating task while pupillometry and electrooculogram signals were recorded simultaneously (example signals in left panel). We then examined how each eye signal tracked the acoustic speech signal, that is, the speech envelope and its first derivative extracted from the waveform (example signals in right panel). (B) Modulation spectrum of both talks. Light and dark green lines indicate average values across 6-s segments for Talk 1 and Talk 2 respectively, with shaded areas representing standard error of the mean. (C) Raincloud plots showing mean arousal and valence ratings across speech chunks for Talk 1 and Talk 2 (*N* = 41). Each point represents an individual participant’s mean rating; half-violin plots illustrate the kernel density of these means. Light grey lines connect each participant’s mean emotion ratings between talks, emphasising within-subject changes. (D) All eye signals track the acoustic speech signal at slow frequencies. Panels show significant speech tracking for pupil, horizontal EOG (hEOG), and vertical EOG (vEOG) from left to right. Talk 1 is represented with a light green line and Talk 2 with a dark green line. The y-axis is logarithmically scaled to emphasise differences across lower frequencies. t-values are derived from comparing z-scored mutual information (MI) values against zero. Light and dark green bars below the x-axis indicate frequency ranges where cluster-based permutation tests revealed significant effects for Talk 1 and Talk 2, respectively.

### EOG data acquisition and preprocessing

EOG data was acquired using a BioSemi ActiveTwo system with a sampling rate of 512 Hz. We used EOG to quantify eye movements, rather than eye-tracker–based gaze measures, because EOG signals are straightforward to obtain without additional calibration demands and provide a meaningful representation of blink-related activity that contributes to the continuous signal. Four Ag/AgCl electrodes were used to capture horizontal and vertical eye movements: two electrodes were placed at the outer canthi of both eyes to measure horizontal EOG (hEOG), and two were placed above and below the left eye to measure vertical EOG (vEOG). A scalp electrode placed at the vertex served as the reference. Electrode signals were visually inspected during recording to ensure data quality. The absolute offset value was kept below 20 μV and never exceeded 30 μV.

EOG data were pre-processed using custom MATLAB scripts (MATLAB 2023a, The MathWorks Inc.) with the FieldTrip toolbox (Oostenveld et al., 2011). Vertical EOG was calculated as the difference between electrodes above and below the right eye, while horizontal EOG was calculated as the difference between electrodes placed at the left and right temples. The continuous EOG recordings were segmented into epochs corresponding to the exact duration of each speech chunk. The segmented EOG signals were then bandpass filtered between 1 Hz and 8 Hz using a fourth-order Butterworth filter (applied forward and reverse) and downsampled to 250 Hz.

### Pupil data acquisition and preprocessing

The pupil signal was recorded using an EyeLink II head-mounted eye-tracking system (SR Research, Osgoode, ON, Canada) for the first cohort of participants (*N* = 23). As several participants reported discomfort with the headset, the remaining participants (*N* = 24) were tested using a desktop-mounted EyeLink 1000 Plus system (SR Research Ltd., Mississauga, ON, Canada) equipped with a chin rest (SR Research Head Support) to stabilise head position. In both setups, pupil signal was continuously recorded from the right eye at a sampling rate of 250 Hz. Before each talk, a standard five-point calibration and validation procedure was performed to ensure accurate pupil measurements. The recording room was dimly lit, and both the fixation circle and background were kept at equal luminance levels to minimise luminance-driven changes in pupil diameter. We conducted a control analysis to assess whether eye-tracking device type influenced pupil–speech mutual information values. Including device type as an additional fixed effect neither improved model fit nor was it a significant predictor of pupil–speech MI values (*χ*²(1) = 1.20, *p* = .27). The patterns of emotion-related effects remained robust, except that the main effect of listeners’ mood arousal was no longer statistically significant after accounting for device type. Overall, the results of the control analysis indicate that the observed findings were not driven by device-related variability (see Supplementary Materials, Table S1).

Pupil data were pre-processed using custom MATLAB scripts (available at https://github.com/anne-urai/pupil_preprocessing_tutorial; as used in Pfeffer et al., 2022; Urai et al., 2017) in combination with functions from the FieldTrip toolbox (Oostenveld et al., 2011). Pupil area values were converted to pupil diameter. Blinks were automatically detected and linearly interpolated, and small eye-tracker artifacts such as spikes and jumps were identified and corrected using the same procedure. We deviated from the preregistration by interpolating blink-related activity in the pupil signal rather than excluding blink windows, as the analysis pipeline required a continuous signal without missing data. Following blink interpolation, the pupil data were band-pass filtered between 1 and 20 Hz using a second-order zero-phase Butterworth filter (applied forward and reverse) to remove slow drifts. To ensure precise temporal alignment between eye-movement and pupil signals, a cross-correlation between z-scored pupil and vertical EOG signals was computed to identify temporal offset. The identified peaks were visually inspected for accuracy. Finally, the pupil and eye movement data were segmented into trials corresponding to the stimulus chunks.

### Speech spectral characteristics and envelope extraction

To compare the spectral characteristics of the two talks, we computed the modulation spectrum for both talks by segmenting speech excerpts into 6-s chunks (Ding et al., 2017; Keitel et al., 2025). Modulation spectra were highly similar, including the 0-12 Hz frequency range relevant to our analysis (Figure 2B). We extracted the wideband speech envelope and its first derivative for each speech chunk. The speech waveform was filtered into eight frequency bands ranging from 100 to 8000 Hz using third-order, forward and reverse Butterworth filters. These bands were spaced equidistantly according to the cochlear frequency scale (Smith et al., 2002). As all recordings were in stereo format, extraction was performed separately for each channel. Within each channel, the filtered signals for each frequency band were Hilbert-transformed, and the magnitude of the signal was extracted to compute the envelope. The resulting envelopes across all eight frequency bands were then averaged for each channel. Finally, the envelopes from both channels were averaged to generate a single wideband envelope per speech chunk. Last, the speech signals were downsampled to 250 Hz.

### Mutual information analysis

We quantified the correspondence between continuous eye and speech signals using a mutual information (MI) framework (Chalas et al., 2022; Ince et al., 2017; Keitel et al., 2017). This approach robustly captures non-linear dependencies between time series using a Gaussian copula-based instead of a binning approach (Ince et al., 2017).

MI (in bits) between the eye signals (continuous pupil dilation, hEOG, and vEOG) and the speech signals (envelope and its derivative) was computed with all signals transformed in the frequency domain (using a continuous wavelet transformation for 46 logarithmically spaced frequencies between 0.5 and 12 Hz) (Chalas et al., 2022). MI values were estimated for 39 lags between 100 ms and 2400 ms (in 60 ms steps), as it is not possible to estimate a single optimal lag per individual for pupil dilation and eye movements. This is because eye activity reflects several cognitive (speech-related and emotion-related) processes, which all follow different temporal dynamics (e.g., Madsen & Parra, 2024). MI values were computed separately for each participant, speech chunk, and eye signal.

Although our preregistration proposed examining MI across frequency bands linked to linguistic units (syllables, words, phrases, sentences), we anchored the analyses to the speech envelope and its first derivative, which offer a more assumption-free representation of speech dynamics.

### Statistical analysis

We first tested whether ocular speech tracking (for pupil dilation, vertical and horizontal EOG) was present when compared against chance-level (i.e., whether MI values for speech-eye correspondence were above chance level). To this end, observed MI values were first normalised using surrogate MI values. This surrogate data was created by randomly shuffling 3-second segments of the speech envelope and derivative (Keitel, Pelofi, et al., 2025). This way, any temporal correspondence between the speech and eye signals was removed. Then, we estimated surrogate MI values for shuffled envelope signals and eye signals. In total, we computed 100 surrogate MI values for each participant, speech chunk, eye signal, and frequency. This was done for 5 equally spaced lags between 100 and 2260 ms, covering the full temporal range of lags. We did not compute surrogate data for all 39 lags as this would have been too computationally intensive. MI values across the five chosen lags were averaged, and surrogate MI was used to z-score observed MI values, which was then compared against zero using a dependent-samples t-tests for all frequencies (Chalas et al., 2022). We used a cluster-based permutation test with a Monte Carlo randomisation procedure (1000 permutations) to correct for multiple comparisons across frequencies (Maris & Oostenveld, 2007). Clusters were identified along the frequency dimension using a cluster-defining threshold of *α* = 0.05 (as well as a cluster-level alpha of *α* = 0.05), and significance was determined based on the 95th percentile of the cluster-level null distribution, controlling the family-wise error rate.

To assess the relationship between emotion-related factors and ocular speech tracking, linear mixed-effects models were fitted separately to each eye signal. The outcome variable was the mutual information between the speech and eye signals. Fixed effects were emotion ratings (i.e. speech arousal and speech valence) and participants’ mood scores (i.e. mood arousal and mood valence). To test whether the relationship between speech emotions and ocular tracking depended on participants’ current mood state, we also included the interaction between emotion ratings and mood scores. We further entered chunk order (to account for changes in tracking over time), talk ID, participant’s empathy scores, and two acoustic predictors (pitch and loudness) into the models as fixed factors. Following the maximal random effects approach (Barr, 2013), we included random intercepts for participants, speech chunks, and speech-eye lags to account for dependencies introduced by the repeated-measures design. Speech-eye lags were included as random factor to get an initial overview of effects that were related to speech tracking (“full model”). All continuous variables were *z*-scored.

To estimate the temporal evolution of effects across the range of lags, we fitted separate models for each individual speech-eye lag. This was identical to the model structure above, except that lag was not included as random factor. The resulting *t*-statistic curves are shown in Supplementary Materials (*Figure S2*).

Overall model fit was evaluated by comparing the full model to a null model containing only random intercepts. For each fixed effect, p-values were obtained using Satterthwaite’s approximation for degrees of freedom (*lmerTest*; Kuznetsova et al., 2017). To control for multiple comparisons across predictors, false discovery rate (FDR) correction was applied to the resulting p-values within each model (Benjamini & Hochberg, 1995). This correction was performed separately for each dependent measure. Diagnostic plots did not reveal any obvious deviations from the assumptions of multilevel analysis (linearity, homogeneity of variance, multicollinearity, and normality of residuals). Models were estimated using restricted maximum likelihood for model comparison purposes, and the significance level was set at *α* = 0.05.

Note that we initially included both linear and quadratic valence terms in the models to capture potential nonlinear effects of valence. These models revealed several main and interaction effects related to quadratic valence. However, the inclusion of the quadratic term introduced complex interaction patterns, obscuring a clear interpretation. For clarity and parsimony, we opted to report the model with only linear component of valence in the main text. Full results from the quadratic models are provided in the Supplementary Materials.

## Results

### Ocular tracking of speech rhythm

We first examined whether participants’ ocular signals tracked the acoustic speech envelope and its first derivative significantly above chance levels using dependent-samples t-tests with cluster-based permutation to control for multiple comparisons across frequencies (see *Figure 2D*). For pupil dilation, this revealed a significant positive cluster for the first talk between 0.53 Hz and 3.45 Hz (cluster *t*_sum_ = 164.34, Cohen’s d_Peak_ = 3.31, *p* = .001) and for the second talk between 0.53 Hz and 6.43 Hz (cluster *t*_sum_ = 233.62, Cohen’s d_Peak_ = 3.12, *p* = .001). For hEOG, a positive cluster emerged between 0.53 Hz and 1.85 Hz (cluster *t*_sum_ = 152.05, Cohen’s d_Peak_ = 4.57, *p* = .001) for the first talk and between 0.53 and 3.69 Hz (cluster *t*_sum_ = 206.56, Cohen’s d_Peak_ = 5.60, *p* = .001) for the second talk. For vEOG, we found a significant positive cluster between 0.53 Hz and 6.89 Hz (cluster *t*_sum_ = 194.12, Cohen’s d_Peak_ = 4.34, *p* = .001) for the first talk, and between 0.53 Hz and 11.20 Hz (cluster *t*_sum_ = 218.02, Cohen’s d_Peak_ = 3.05, *p* = .001) for the second talk. These findings suggest that all eye signals systematically tracked slow fluctuations in the acoustic speech signal.

### Predictors of pupil speech tracking

We evaluated the relationship between pupil speech tracking and our predictors (speech arousal, speech valence, mood arousal, mood valence, trait empathy, pitch, loudness, speech chunk order, and talk ID) with a linear mixed-effects model. The full model included all used speech-eye lags as random factors to assess the overall role of predictors. Additionally, we computed single-lag models to assess the temporal evolution of each predictor (see supplementary materials *Figure S2*).

We observed a positive main effect of *speech arousal* (*β* = 0.02, *p*_FDR_ < .001) in the main model, suggesting stronger pupil tracking for high-arousal speech than low-arousal speech. Single-lag models revealed peak effects of arousal at 1780 ms. We also observed a main effect of *mood arousal* (*β* = 0.04, *p*_FDR_ = .04), with individuals in high-arousal mood tracking speech more strongly than those in low-arousal mood. No significant main effects were observed for the remaining predictors (see Table 2).

**Table 2.** Summary table for main model predicting pupil-speech MI values. LL = Lower limit, UL = Upper limit, A = Arousal dimension, V = Valence dimension. Fixed-effect estimates (β) are reported with confidence intervals (95% CI) computed using the Wald method. *p* values were false discovery rate (FDR) corrected. Significant predictors are highlighted in bold. The model included random intercepts for participants, speech chunks, and speech-eye lags. *\*p* <.05*, **p* <.001.

| Predictors | Estimates | 95% CI | | $p_{\text{uncorrected}}$ | $p_{\text{FDR}}$ |
| --- | --- | --- | --- | --- | --- |
|  |  | LL | UL |  |  |
| Intercept | 0.12 | -0.65 | 0.90 | .752 | .806 |
| **Arousal** | 0.02 | 0.01 | 0.02 | **<.001\*\*** | **<.001\*\*** |
| Valence | -0.01 | -0.01 | -0.001 | .035* | .065 |
| **Mood (A)** | 0.04 | 0.01 | 0.07 | .009* | .038* |
| Mood (V) | 0.01 | -0.02 | 0.04 | .400 | .552 |
| Time-in-talk | 0.03 | -0.08 | 0.14 | .609 | .763 |
| Trait empathy | -0.03 | -0.06 | -0.01 | .015* | .050 |
| Pitch | 0.11 | -0.04 | 0.26 | .165 | .253 |
| Loudness | -0.02 | -0.24 | 0.20 | .837 | .838 |
| Talk | -0.09 | -0.61 | 0.44 | .745 | .806 |
| **Arousal × Valence** | 0.02 | 0.02 | 0.03 | **<.001\*\*** | **<.001\*\*** |
| **Arousal × Mood (A)** | 0.015 | 0.01 | 0.02 | **<.001\*\*** | **<.001\*\*** |
| Arousal × Mood (V) | -0.004 | -0.01 | 0.001 | .089 | .148 |
| **Valence × Mood (A)** | -0.01 | -0.015 | -0.01 | **<.001\*\*** | **<.001\*\*** |
| Valence × Mood (V) | -0.005 | -0.01 | -0.000 | .034* | .065 |

The full model also revealed several interaction effects. First, we found a two-way interaction effect of *speech arousal* × *speech valence*. This suggests that when speech was positive in valence, high-arousal speech was tracked more strongly than low-arousal speech (*β* = 0.02, *p*_FDR_ < .001; peak effect observed at 880 ms in the single-lag models). In contrast, when speech was negative in valence, this relationship reversed, with low-arousal speech being tracked more strongly (see *Figure 3A*).

**Fig. 3.**
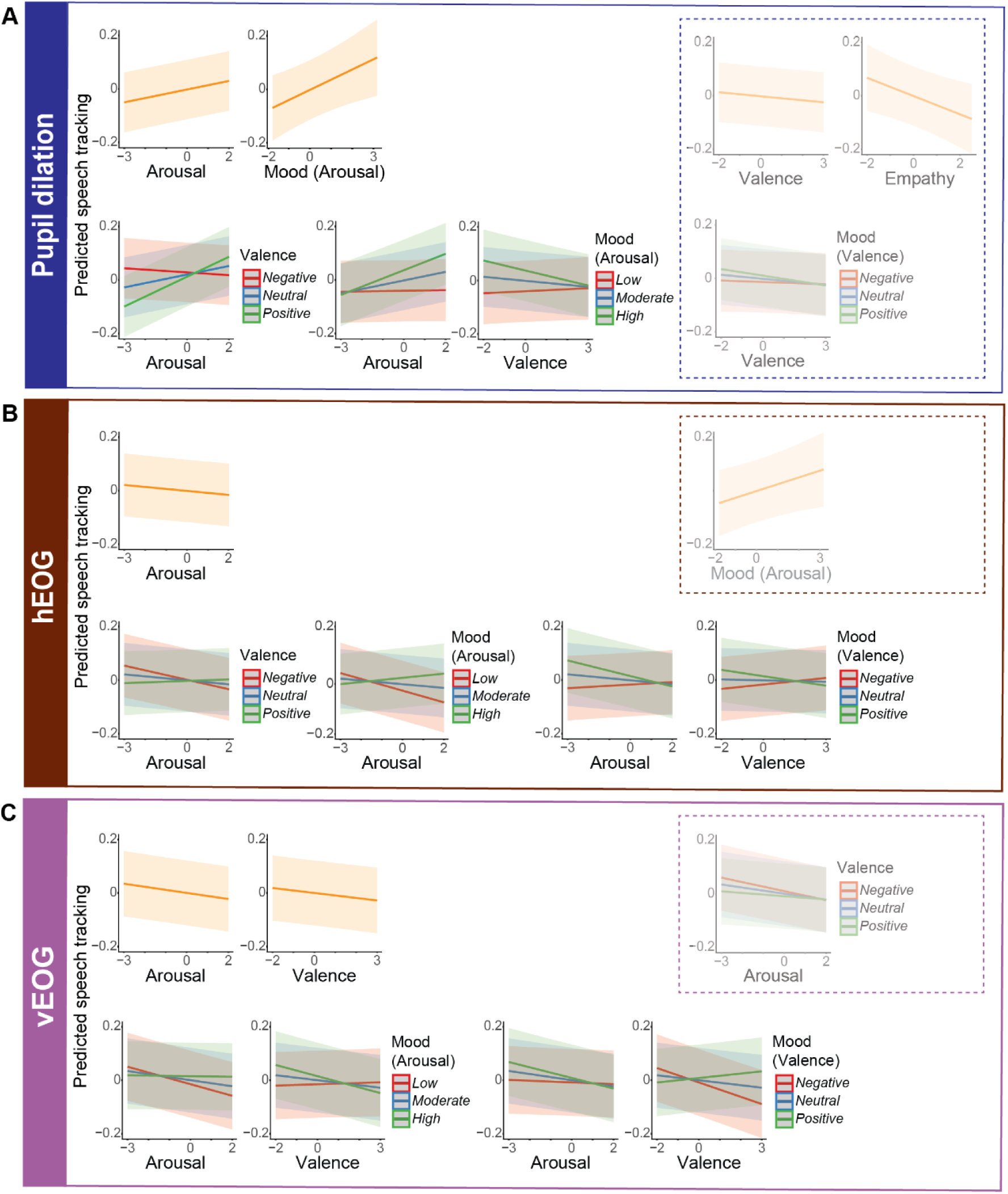
Relationship between ocular tracking of the speech signal and emotion-related predictors. The figures were derived from separate linear mixed-effects models with pupil–envelope MI, hEOG–envelope MI, and vEOG–envelope MI as outcome variables (top to bottom). Each plot displays the predicted MI values from the model (y-axis) as a function of the corresponding predictor (x-axis), with shaded areas representing 95% confidence intervals. Effects shown within the dashed outline were not significant after FDR correction. (A) For pupil dilation, the model revealed significant main effects of arousal and mood arousal on pupil tracking of speech signal. Interaction effects of arousal × valence, arousal × mood arousal, and valence × mood arousal also predicted pupil tracking of speech. (B) For horizontal EOG, a significant main effect of arousal emerged, along with interaction effects of arousal × valence, arousal × mood arousal, arousal × mood valence and, valence × mood valence. (C) For vertical EOG, the main effect of arousal and valence, as well as the interaction effects of arousal × mood arousal, valence × mood arousal, arousal × mood valence, valence × mood valence predicted envelope tracking. Overall, ocular tracking of speech was influenced by both stimulus-driven and listener-dependent emotional factors.

Second, we found interaction effects between speech emotion (i.e. arousal or valence ratings) and individuals’ own mood (both arousal and valence dimensions). When participants were in a high-arousal mood, they tracked high-arousing speech stronger, while this relationship was absent for people in a low-arousal mood (*arousal × mood arousal*: *β* = 0.015, *p*_FDR_ < .001; peak effect observed at 2020 ms in the single-lag models). Additionally, individuals in a high-arousal mood tracked negative speech more strongly than positive speech, whereas those in a low-arousal mood tended to track positive speech more strongly (*speech valence* × *mood arousal*: *β* = −0.01, *p*_FDR_ < .001; peak effect observed at 2260 ms in the single-lag models). Regarding *mood valence*, while we did not observe a significant effect for its interaction with speech arousal; however, *speech valence* × *mood valence* predicted pupil speech tracking. Yet, this effect did not survive correction for multiple comparisons (p_uncorrected_ = .04; *p*_FDR_ =.06)

### Predictors of hEOG speech tracking

The full model with hEOG-speech tracking as outcome variable revealed that only the main effect of *arousal* was significant (*β* = −0.007, *p*_FDR_ = .031; peak effect observed at 1600 ms in the single-lag models), suggesting that horizontal eye movements tracked low-arousal speech stronger than high-arousal speech. *Mood arousal* also predicted hEOG-speech coupling, however its effect did not survive after FDR correction. None of the remaining predictors showed a significant main effect (see Table 3).

**Table 3.** Summary table for the mixed effects model predicting hEOG-speech MI values. A = Arousal dimension, V = Valence dimension. Fixed-effect estimates (β) are reported with confidence intervals (95% CI) computed using the Wald method. *p* values were false discovery rate (FDR) corrected. Significant predictors are highlighted in bold. The model included random intercepts for participants speech chunks and speech-eye lags. *\*p <.05 **p<.001*.

| Predictors | Estimates | 95% CI | | $p_{uncorrected}$ | $p_{FDR}$ |
| --- | --- | --- | --- | --- | --- |
|  |  | LL | UL |  |  |
| Intercept | 0.31 | -0.49 | 1.11 | .446 | .611 |
| **Arousal** | -0.01 | -0.01 | -0.002 | .011* | **.033\*** |
| Valence | -0.001 | -0.01 | 0.005 | .700 | .731 |
| Mood (A) | 0.02 | 0.01 | 0.05 | .038* | .112 |
| Mood (V) | 0.02 | -0.01 | 0.04 | .185 | .361 |
| Time-in-talk | 0.02 | -0.09 | 0.13 | .731 | .731 |
| Trait empathy | -0.01 | -0.03 | -0.01 | .502 | .632 |
| Pitch | 0.07 | -0.09 | 0.23 | .382 | .611 |
| Loudness | 0.06 | -0.17 | 0.29 | .605 | .699 |
| Talk | -0.21 | -0.77 | 0.33 | .438 | .611 |
| **Arousal × Valence** | 0.01 | 0.005 | 0.01 | <.001** | **<.001\*\*** |
| **Arousal × Mood (A)** | 0.02 | 0.01 | 0.02 | <.001** | **<.001\*\*** |
| **Arousal × Mood (V)** | -0.01 | -0.02 | -0.01 | <.001** | **<.001\*\*** |
| Valence × Mood (A) | -0.004 | -0.01 | 0.001 | .086 | .185 |
| **Valence × Mood (V)** | -0.01 | -0.01 | -0.01 | <.001** | **<.001\*\*** |

The interaction between *speech arousal* and *speech valence* was significant (*β* = 0.01, *p*_FDR_ < .001; peak effect observed at 340 ms in the single-lag models). Specifically, hEOG tracking was stronger for low-arousal speech when speech was perceived as more negative, whereas high arousal speech was tracked more strongly when speech was perceived as more positive (see *Figure 3B*).

The model further revealed significant interaction effects between participants’ mood and emotional characteristics of speech. When participants were in a low-arousal mood, low-arousal speech was tracked more strongly than high-arousal speech, in contrast, when participants were in a high-arousal mood, tracking was stronger for high-arousal than for low-arousal speech (*speech arousal* × *mood arousal*: *β* = 0.02, *p*_FDR_ < 0.001; peak effect observed at 820 ms in the single-lag models). Participants in a positive mood tracked low-arousal speech more strongly than high-arousal speech, whereas those in a negative mood exhibited the opposite pattern (*speech arousal* × *mood valence*: *β* = −0.01, *p*_FDR_ < .001; peak effect observed at 1300 ms in the single-lag models). In a positive mood, negative speech was tracked more strongly than positive speech, while in a negative mood, positive speech was tracked more strongly (*speech valence* × *mood valence*: *β* = −0.01, *p*_FDR_ < .001; peak effect observed at 640 ms in the single-lag models).

### Predictors of vEOG speech tracking

The full model predicting vEOG speech tracking revealed a significant main effect of *speech arousal* (*β* = −0.01, *p*_FDR_ < .001; peak effect observed at 1480 ms in the single-lag models), indicating stronger tracking for low-than high-arousal speech. The model also revealed a significant main effect of *speech valence* (*β* = −0.01, *p*_FDR_ = .01; peak effect observed at 100 ms in the single-lag models), with stronger tracking of negative speech than positive speech (see Table 4).

**Table 4.** Summary table for the mixed effects model predicting vEOG-speech MI values. A = Arousal dimension V = Valence dimension. Fixed-effect estimates (β) are reported with confidence intervals (95% CI) computed using the Wald method. p values were false discovery rate (FDR) corrected. Significant predictors are highlighted in bold. The model included random intercepts for participants, speech chunks and speech-eye lags. *p <.05 **p<.001.

| Predictors | Estimates | 95% CI | | $p_{\text{uncorrected}}$ | $p_{\text{FDR}}$ |
| --- | --- | --- | --- | --- | --- |
|  |  | LL | UL |  |  |
| Intercept | 0.20 | -0.63 | 1.03 | .640 | .740 |
| **Arousal** | -0.01 | -0.02 | -0.01 | <.001** | <.001** |
| **Valence** | -0.01 | -0.02 | -0.004 | .003* | .007* |
| Mood (A) | 0.02 | -0.02 | 0.05 | .355 | .601 |
| Mood (V) | 0.01 | -0.02 | 0.04 | .592 | .740 |
| Time-in-talk | 0.02 | -0.10 | 0.14 | .772 | .828 |
| Trait empathy | -0.01 | -0.04 | 0.02 | .594 | .740 |
| Pitch | 0.11 | -0.06 | 0.27 | .217 | .411 |
| Loudness | -0.01 | -0.25 | 0.22 | .909 | .909 |
| Talk | -0.14 | -0.70 | 0.43 | .636 | .740 |
| Arousal × Valence | 0.01 | 0.001 | 0.01 | .025* | .054 |
| **Arousal × Mood (A)** | 0.01 | 0.01 | 0.01 | <.001** | <.001** |
| **Arousal × Mood (V)** | -0.01 | -0.01 | 0.004 | <.001** | .003* |
| **Valence × Mood (A)** | -0.01 | -0.02 | -0.01 | <.001** | <.001** |
| **Valence × Mood (V)** | 0.02 | 0.01 | 0.02 | <.001** | <.001** |

Interaction effect between *speech arousal* and *speech valence* predicted pupil speech tracking. However, this effect did not survive correction for multiple comparisons (p_uncorrected_ = .02; *p*_FDR_ =.054).

Significant interactions were observed between participants’ mood and speech emotions. For mood arousal, participants in a low-arousal mood showed stronger tracking of low-arousal speech compared to high-arousal speech, whereas this pattern was not observed for those in a high-arousal mood (*speech arousal* x *mood arousal*: *β* = 0.01, *p*_FDR_ < 0.001; peak effect observed at 1540 ms in the single-lag models). Participants in a low-arousal mood also tracked positive speech stronger than negative speech, in contrast, participants in a high-arousal mood tracked negative speech stronger than positive speech (*speech valence* x *mood arousal: β* = −0.01, *p*_FDR_ < 0.001; peak effect observed at 880 ms in the single-lag models).

When participants were in a negative mood, tracking of negative speech was stronger than that of positive speech, whereas this pattern reversed for those in a positive mood, who showed stronger tracking of positive than negative speech (*speech valence* x *mood valence*: *β* = 0.02, *p*_FDR_ < .001; peak effect observed at 2020 ms in the single-lag models). Moreover, in a positive mood, the tracking of low-arousal speech was stronger than high-arousal speech, with this relationship is absent for negative mood (*speech arousal* x *mood valence*: *β* = −0.01, *p*_FDR_ = .003; peak effect observed at 700 ms in the single-lag models).

### Differences in emotion ratings between talks

We tested whether emotional responses differed across the two TED talks by fitting separate linear mixed-effects models for arousal and valence ratings, with Talk (Talk 1 vs. Talk 2) included as a fixed effect. Consistent with the results of Study 1, arousal ratings did not differ between the two talks (*p*_FDR_ = .997). In contrast, valence ratings were significantly more positive for Talk 2 compared to Talk 1 (*β* = 0.17, 95% CI [0.14 – 0.20], *p* < .001, *p*_FDR_ < .001). Chunk-level distributions of arousal and valence ratings are shown in *Figure 2C*.

## Discussion

In the present study, we show that ocular dynamics (pupil dilation, horizontal and vertical EOG) track the quasi-rhythmic fluctuations of the speech envelope at low frequencies (0.5 - 12 Hz). Moreover, ocular speech tracking is sensitive to emotion-related factors, including emotional qualities embedded in the speech (as reflected by subjective ratings) and listener-dependent states, such as mood, and trait empathy.

### Pupil and eye movements significantly track speech at slow frequencies

Listeners showed reliable ocular tracking of the speech signal in our study, suggesting that bodily rhythms, much like neural rhythms, follow the temporal dynamics of continuous speech. Ocular speech tracking was observed at slow frequencies between 0.5 Hz and 12 Hz, with each eye-related signal showing somewhat different its frequency ranges. The specific frequency bands exhibiting significant tracking further varied across talks, which may be related to acoustic characteristics.

Previous studies have shown that vertical EOG activity, largely reflecting eye blinks, tracks higher-level linguistic structures, whereas horizontal EOG and pupil signals were not found to couple with speech features (Jin et al., 2018). More recent gaze-based research, however, has demonstrated that both vertical and horizontal eye movements track the speech signal, with different tracking patterns (tracking of short sentences: Gehmacher et al., 2024; tracking of continuous speech: Schubert et al., 2025). Regarding pupil activity, in contrast to the findings by Jin et al. (2018), more recent work has shown that pupil responses align with word onsets and the speech envelope (Madsen & Parra, 2024). It is important to acknowledge that direct comparisons between these limited numbers of studies are challenging, given the variations in study designs, the specific speech features examined, and the methods to quantify speech tracking. Our findings support the presence of a coupling between eye-related physiological signals and the acoustic dynamics of continuous speech, providing further evidence for the precise temporal characteristics of this relationship. Importantly, we interpolated blinks and removed the canonical responses to saccades and blinks from the pupil signals (Lehmann & Corneil, 2016; Urai et al., 2017; Wang & Munoz, 2015), thereby, ensuring that the observed pupil–speech coupling reflects processes distinct from eye movement related activity.

We are still far from understanding the underlying mechanisms and functional relevance of ocular speech tracking. One proposed explanation is that it may serve as an active sensing mechanism, whereby motor actions are engaged to optimise perception, a notion suggested by predictive coding accounts (Friston, 2010; Schubert et al., 2025). The involvement of motor systems in speech perception has long been established (Lane, 1965; Scott et al., 2009). The aforementioned growing body of literature adopting speech tracking measures extends this view by providing direct evidence for the involvement of eye movements in speech processing. Specifically, by aligning sensory systems with the incoming speech stream, eye movements could increase sensory gain and reduce uncertainty in dynamic listening environments (Schubert et al., 2025). While this perspective offers a promising framework, substantiating it will require systematic empirical work that integrates ocular, neural, and behavioural measures of speech processing.

### Subjective emotion ratings predict ocular tracking of speech

In line with our expectations, listeners’ subjective emotion ratings influenced the strength of ocular speech tracking, with distinct patterns across signals. The effect of valence followed a similar pattern for pupil and vertical tracking of speech: negative speech was associated with stronger speech tracking than positive speech. Note the effect of speech valence on pupil envelope tracking did not survive after FDR correction. However, the quadratic model (see supplemental material) revealed a robust effect of the polynomial term of speech valence, suggesting stronger pupil tracking of neutral than positively or negatively valenced speech.

Valence is one of the most fundamental dimensions of emotional experience. A large body of findings show that both positive and negative linguistic stimuli hold processing advantages over neutral stimuli across diverse experimental designs (Kim & Sumner, 2017; Kousta et al., 2009; Schirmer et al., 2005). However, it remains unclear whether stimuli at opposite ends of the continuum confer distinct advantages, and if so, in favour of which. Evidence suggests a processing advantage for positive words over neutral and negative words in visual lexical decision tasks (Bayer & Schacht, 2014; Hofmann et al., 2009; Kauschke et al., 2019). A recent Bayesian meta-analysis also found a robust facilitative effect of positive valence on lexical decision times; effects of negative valence, however, were only observed occasionally, specifically when negative words were perceived as highly intense (Ferré et al., 2025). Some evidence suggests that valence effects appear to be non-linear, with a U-shaped pattern such that both positive and negative words are recognised faster than neutral words in an auditory lexical decision task (Gao et al., 2022).

Emotional stimuli may bias attentional mechanisms to facilitate sensory processing (Pessoa, 2008; Vuilleumier, 2005). In the auditory domain, electrophysiological studies suggest an enhanced spatial attention to negatively valence stimuli (i.e. threatening prosody), compared to positive valence (i.e. happy prosody) and neutral stimuli (Burra et al., 2019). Behavioural evidence also suggests that temporal attention biases for angry prosody, in comparison to neutral prosody (Guex et al., 2022). Our findings may be interpreted within this framework: negative speech may have captured listeners’ attention more effectively, leading to stronger alignment between ocular responses and the speech signal. While this interpretation aligns with previous evidence suggesting that increased attentional engagement enhances the strength of speech tracking (Gehmacher et al., 2024; Schubert et al., 2025), which should be approached with caution given that speech stimuli used in the present study do not convey the same level of threat as the stimuli typically employed in the prior studies. Moreover, our study does not provide direct evidence for the role of attention in this relationship, and future studies are necessary to explore the interplay between emotion, attention, and ocular tracking in naturalistic paradigms.

The effects of arousal revealed distinct patterns for pupil and eye movement tracking of speech. For pupil responses, higher arousal was associated with stronger speech tracking, whereas both horizontal and vertical EOG signals tracked low-arousal speech stronger. Moreover, we observed a significant interaction between arousal and valence in predicting pupil and horizontal EOG tracking. The main effect pattern of arousal was repeated for the pupil tracking only in response to positive speech, which reversed for negative speech segments (i.e. low arousal led to stronger tracking in response to negative speech). For horizontal EOG, higher arousal enhanced tracking only when the speech was perceived as more positive. Overall, these results suggest that pupil and eye movements show distinct responses to high arousal speech. The sensitivity of pupil responses to arousal is well-established (Bradley et al., 2008; Pfeffer et al., 2022), and there is some evidence for an enhanced pupil response to emotional speech (Jürgens et al., 2018). Yet, pupil responses also index other cognitive processes, such as attention (Koelewijn et al., 2014), or listening effort (Zekveld et al., 2010). While these processes cannot be disentangled in the present paradigm, future work should adopt experimental designs that isolate the contributions of attention, effort, and arousal more directly.

Our lag-based models further revealed that the influence of emotion-related predictors on ocular speech tracking unfolds over distinct temporal scales (see Supplementary Materials, Figure S3). Specifically, we observed an early effect of valence on vEOG speech tracking that peaked at around 160 ms and 100 ms, respectively, whereas the main effect of arousal consistently peaked at later lags (∼1480-1780 ms) across all eye signals. The interaction between speech arousal and mood-related predictors also predominantly peaked at later time windows, typically between 1400 ms and 2200 ms. This temporal dissociation suggests that listeners’ judgements about stimulus valence (positivity or negativity) precede their arousal judgements in ocular speech tracking. Previous work has demonstrated that different cognitive and emotional processes are reflected at distinct temporal stages of the pupil response, spanning early and late components (Madsen & Parra, 2024). Extending these findings, here we show that both horizontal and vertical EOG signals exhibit similar multi-stage temporal dynamics during emotional speech tracking, suggesting that eye movements might also capture a mixture of fast and slower processes during continuous speech perception.

To our knowledge, no previous study has directly examined how emotion modulates the tracking of continuous speech, whether at the physiological or cortical level. However, related evidence from research on infant-directed speech may offer an intriguing point of comparison. Infant-directed speech has been linked to enhanced cortical tracking in infant listeners with distinct synchronisation patterns relative to adult-directed speech (Attaheri et al., 2022; Kalashnikova et al., 2018; Menn et al., 2022). Since infant-directed speech shares similar acoustic features with emotionally expressive speech, particularly its enhanced prosodic stress, it has been considered as an emotional form of speech (Saint-Georges et al., 2013; Trainor et al., 2000). Building on this resemblance, albeit speculatively, our findings may reflect a similar mechanism, whereby emotional speech facilitates temporal alignment to the acoustic envelope of speech in ocular responses.

Another consideration is that the causal direction of this relationship remains unclear While stronger tracking may occur because of perceiving the speech as more emotional, the reverse could also be plausible, where listeners perceive the speech as more emotional because they exhibit stronger tracking. Indeed, enhanced cortical tracking has been proposed as a mechanism underlying emotional induction in the music domain (Trost et al., 2017; Juslin et al., 2024), and empirical support for this account is growing (Ding et al., 2025).

### Participant’s mood modulates the relationships between emotion and speech tracking

The interaction patterns in our main models indicate that listeners’ mood shaped the relationship between speech emotions and envelope tracking, yet the nature of this modulation depended on the specific ocular signal. Across all signals, mood arousal interacted with speech arousal, and mood valence interacted with speech valence.

In high-arousal mood, high-arousal speech was associated with stronger tracking of the speech envelope in both the pupil and hEOG. Note that, for pupil dilation, the main effect of mood arousal also significantly predicted envelope tracking, that is listeners in high arousal mood showed overall strong tracking of speech. In low-arousal mood, low-arousal speech was associated with stronger tracking in both vEOG and hEOG, while this pattern was absent for pupil tracking. Mood arousal occasionally interacted with speech valence; in high-arousal mood, the tracking of negative speech was stronger than that of positive speech in both the pupil and vEOG, whereas this relationship was weaker in low arousal mood.

For the interaction between mood valence and speech valence, the patterns differed across ocular signals, with opposing effects for vEOG and hEOG, and a relatively weaker effect for pupil tracking. For listeners in a negative mood, negative speech was associated with stronger tracking in both the pupil and vEOG, whereas in hEOG positive speech led to stronger tracking than negative speech. In positive mood, pupil and hEOG tracking of negative speech was stronger. In contrast, vEOG showed the opposite pattern, with positive speech being associated with stronger tracking than negative speech. Moreover, significant interactions between mood valence and speech arousal were observed for eye-movement–based tracking. In both vEOG and hEOG, low-arousal speech was associated with stronger tracking in positive mood; in negative mood, this pattern reversed for hEOG, and no relationship between tracking and arousal observed for vEOG.

Many researchers stress the fundamental role of mood in information processing across domains, including language processing (Egidi & Nusbaum, 2012; Van Berkum et al., 2013). There is a general understanding of how positive or negative mood could influence the style of information processing. Specifically, positive mood is thought to promote a broader engagement with stimuli, whereas negative mood is linked with narrower, and more stimulus-oriented processing (Clore & Huntsinger, 2007; Forgas, 2017). A congruency bias has also been suggested, such that stimuli with a valence that matches one’s emotional state receive processing advantages (Bower, 1981). Our results are partially consistent with a congruency bias. In high-arousal mood, high-arousal speech was associated with stronger tracking, and the same held for low-arousal speech in low-arousal mood. However, it was less straightforward for mood valence: while congruency effects were evident in pupil and vEOG tracking, it did not apply to hEOG tracking of the speech envelope. Overall, our findings suggest a complex interaction between the emotional qualities of the speech and the listener’s mood which partly differed across eye signals, highlighting the need for further work to test these relationships in naturalistic listening contexts.

### Conclusion

In conclusion, our first study demonstrated that perceived emotions varied over time within each talk. Our models indicated that speech at both ends of the valence spectrum was associated with higher arousal ratings. Participants in a positive mood provided higher valence ratings, and greater empathy was associated with higher arousal ratings. In our second study, we showed that eye signals, including pupil dilation as well as vertical and horizontal EOG activity, tracked speech at low frequencies. This finding supports the view that ocular responses may act as an active sensing mechanism that contributes to speech processing even in the absence of visual input. We also found that subjective emotion ratings, and their interactions with listeners’ mood, modulated the tracking of speech signals, with distinct patterns for each signal. Together, our findings highlight the relevance of considering both the emotional characteristics of the stimuli and listener-dependent factors, such as transient mood states, in research on speech tracking, which has typically relied on neutral or positive speech materials.

## Acknowledgments

We thank all participants who so generously gave their time and effort to take part in this study. We also thank Efstratios Koukouvinis and Ruta Cinite for their support in data collection. SŞA is funded by the Republic of Türkiye, Ministry of National Education, for her doctoral studies. AK is supported by the Medical Research Council [grant number MR/W02912X/1]. AK and CK are members of the Scottish-EU Critical Oscillations Network (SCONe), funded by the Royal Society of Edinburgh (RSE Saltire Facilitation Network Award to CK and AK, Reference Number 1963). The funders had no involvement in the study protocol, participant recruitment, data analysis, or manuscript preparation.

## Data availability statement

Data and stimuli are publicly available on the OSF (https://osf.io/dsvzh/files/osfstorage).

## Author contributions

S.S.A.: Conceptualisation, methodology, formal analysis, investigation, data curation, writing—original draft, writing—review and editing, visualisation, funding acquisition. C.K.: Software, writing—review & editing, supervision. R.T.: Software, writing—review, A.K.: Conceptualisation, methodology, formal analysis, investigation, software, writing—review and editing, supervision, funding acquisition.

## Supplementary Materials

### 1. Chunk-level distributions of arousal and valence ratings

In Study 1, we showed that participants’ arousal and valence ratings varied across individual speech chunks within each talk. In the main report, we presented raincloud plots summarising the mean participant ratings per talk. Here, we provide chunk-level ridge plots to illustrate how emotion ratings fluctuate across the full sequence of chunks within each talk.

**Figure S1.**
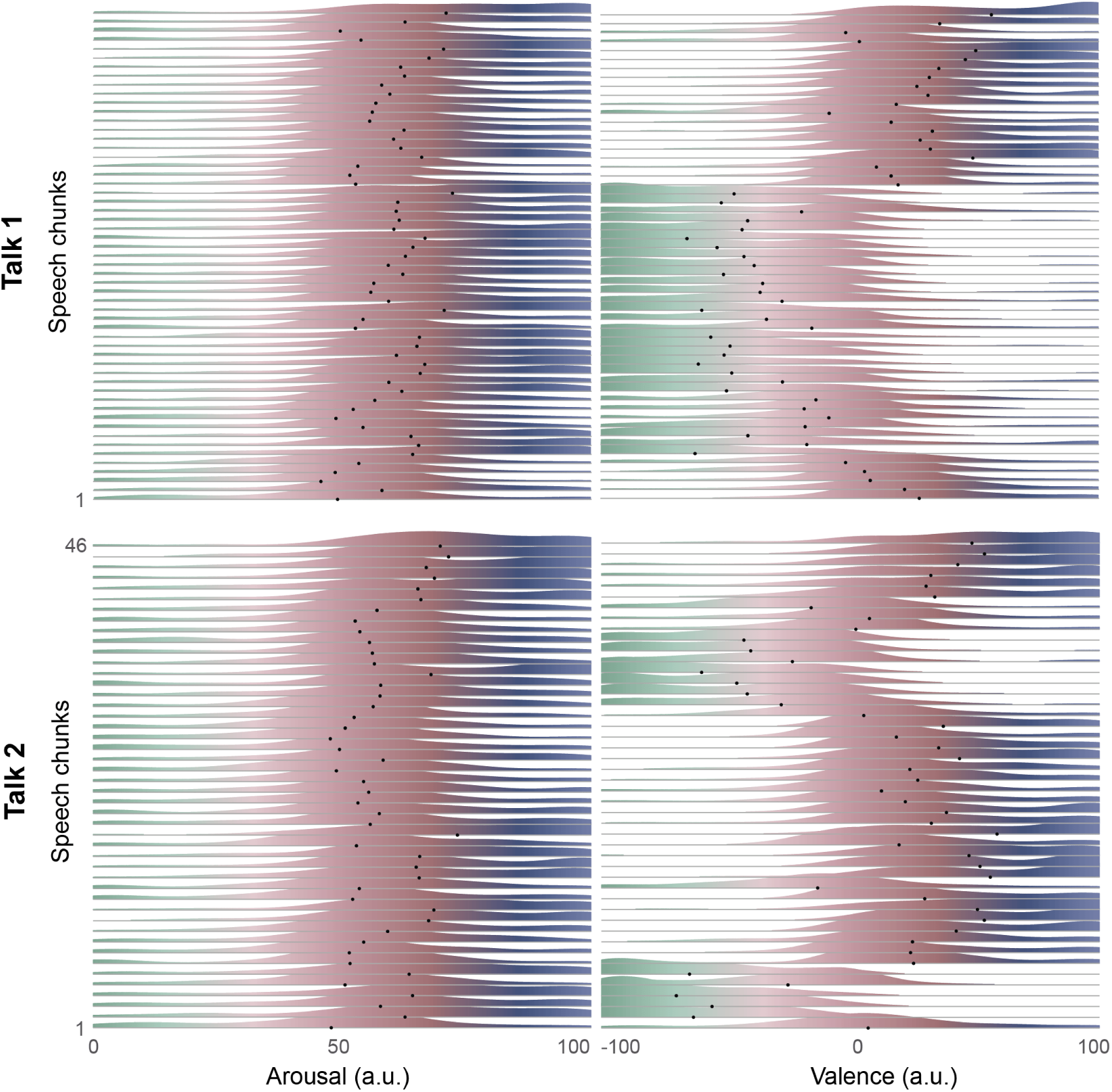
Ridgeplots with distribution of emotion ratings for both talks. Each horizontal line (i.e. y-axis) represents the density of ratings (arousal left, valence right) for a single speech chunk. x axis represent emotional arousal (between 0 and 100) and valence (between −100 and 100) ratings. Colouring reflects the rating level: red indicates mid-range values, blue reflects high arousal or positive valence, and green reflects low arousal or negative valence. Black dots indicate the mean value for each chunk.

Both perceived arousal and valence vary over time, although valence does more so than arousal. Notably, both talks include sections that are perceived as positive and sections that are perceived as negative. This confirms the suitability of the material for using it to study emotion-related ocular tracking responses.

### 2. Device differences for pupil envelope tracking

To assess whether the use of different eye-tracker devices influenced acoustic envelope tracking in the pupil signal or its relationship with emotion-related parameters, we fitted a linear mixed-effects model. This model was identical to the full envelope-tracking models, except that device type was added as a fixed effect. The analysis showed that device type did not predict MI values between the pupil signal and the speech envelope, and the pattern of significant predictors remained mostly unchanged relative to the full model (i.e., the model without device type)(see Table 1). This indicates that hardware differences did not influence our results for pupil-speech coupling.

**Table S1.** Summary table for control model predicting pupil-speech MI values. LL = Lower limit, UL = Upper limit, A = Arousal dimension, V = Valence dimension. Fixed-effect estimates (β) are reported with confidence intervals (95% CI) computed using the Wald method. *p* values were false discovery rate (FDR) corrected. Significant predictors are highlighted in bold. Blue p-values indicate effects that were significant in the full model excluding device as a predictor. The model included random intercepts for participants, speech chunks, and speech-eye lags. *\*p* <.05*, **p* <.001

| Predictors | Estimates | 95% CI | | $p_{uncorrected}$ | $p_{FDR}$ |
| --- | --- | --- | --- | --- | --- |
|  |  | LL | UL |  |  |
| Intercept | 0.07 | -0.70 | 0.85 | .85 | .85 |
| **Arousal** | 0.02 | 0.01 | 0.02 | <.001** | <.001** |
| Valence | -0.01 | -0.01 | -0.001 | .03* | .07 |
| Mood (A) | 0.03 | 0.004 | 0.06 | .03* | .07 |
| Mood (V) | 0.01 | -0.02 | 0.04 | .60 | .75 |
| Time-in-talk | 0.03 | -0.08 | 0.14 | .61 | .75 |
| Trait empathy | -0.03 | -0.06 | -0.01 | .02* | .054 |
| Pitch | 0.11 | -0.04 | 0.26 | .17 | .27 |
| Loudness | -0.02 | -0.24 | 0.20 | .84 | .85 |
| Talk | -0.09 | -0.61 | 0.44 | .75 | .85 |
| Device | 0.03 | -0.02 | 0.09 | .28 | .40 |
| **Arousal × Valence** | 0.02 | 0.02 | 0.03 | <.001** | <.001** |
| **Arousal × Mood (A)** | 0.01 | 0.01 | 0.02 | <.001** | <.001** |
| Arousal × Mood (V) | -0.005 | -0.01 | 0.001 | .09 | .15 |
| **Valence × Mood (A)** | -0.01 | -0.015 | -0.01 | <.001** | <.001** |
| Valence × Mood (V) | -0.005 | -0.01 | -0.000 | .03* | .07 |

### 3. Time-resolved model results

To estimate the temporal evolution of effects across the range of lags, we fitted separate models for each individual speech-eye lag (39 in total, between 100 and 2260 ms). The outcome variable was the mutual information between the speech signal and eye signals. The model structure was identical to those in main model: fixed effects were emotion ratings (i.e. speech arousal and speech valence) and participants’ mood scores (i.e. mood arousal and mood valence), as well as the interaction between emotion ratings and mood scores. We further entered chunk order, participant’s empathy scores, and two acoustic predictors (pitch and loudness) and talks as fixed effects into the models. Random factors included participants and speech chunks. These models allowed us to observe the temporal evolution of effects and to identify the time points at which the peak effects (i.e., the highest t-value) occurred. All continuous variables were *z*-scored.

**Figure S2.**
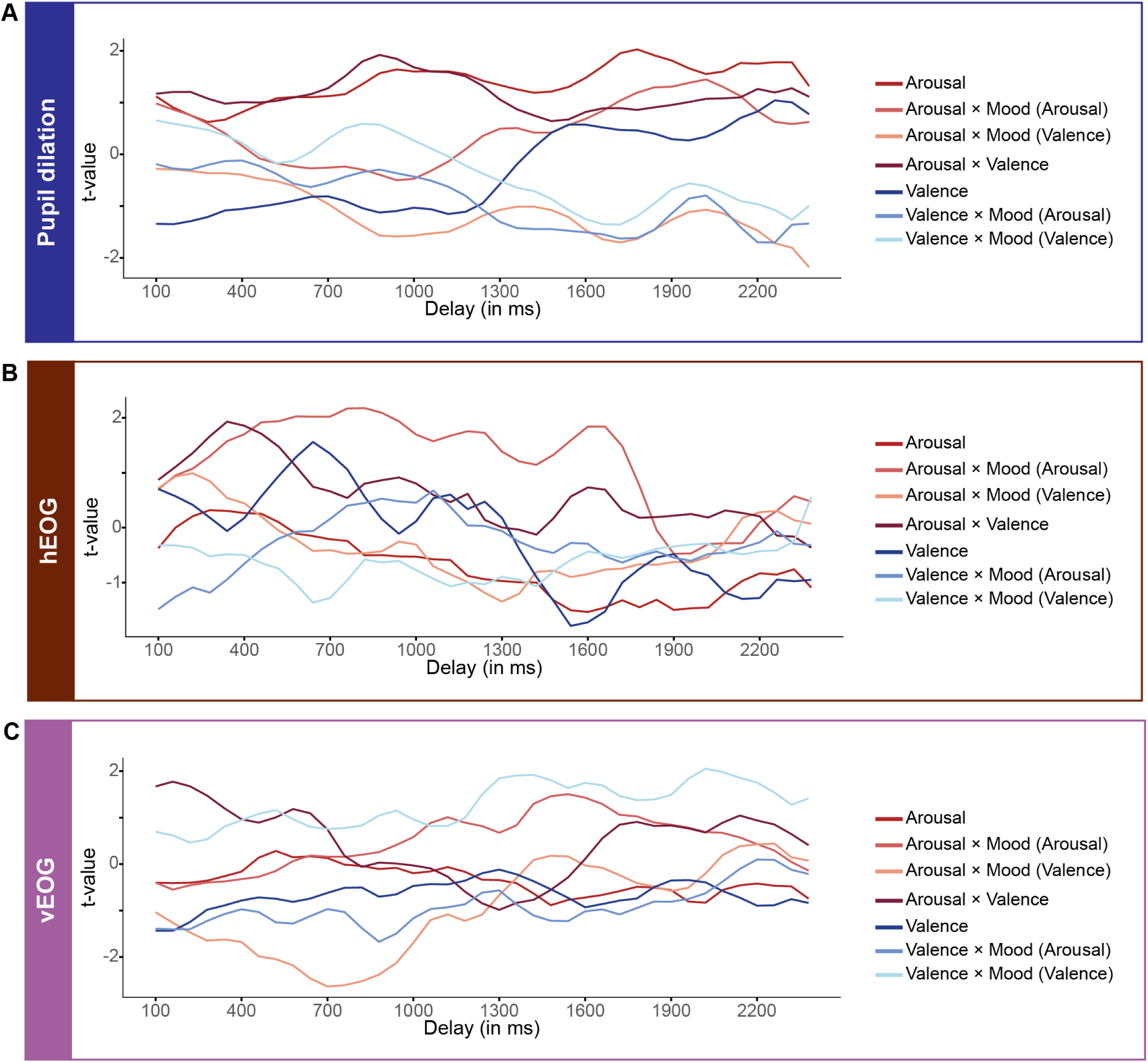
t-statistic curves across lags for each signal. (A) Main model results for pupil tracking of the speech signal revealed a positive, significant effect of speech arousal with its influence on pupil tracking with a peak at approximately 1780 ms. While the effect of speech valence did not significant after FDR correction, this negative effect reached its peak at around 160 ms. The significant interaction between speech arousal and speech valence, which reached its peak at approximately 880 ms. The interaction effects between arousal and mood arousal (peak effect at 2020 ms); valence and mood arousal (peak effect at 2260 ms); valence and mood valence (not significant after FDR; peak at 1720) were also observed. (B) For hEOG tracking of the speech signal, the main effect of arousal was significant (peak effect at 1600 ms). The interaction between speech arousal and speech valence (peak effect at 340 ms); arousal and mood valence (peak effect at 1300 ms); valence and mood valence (peak effect at 640 ms); and between arousal and mood arousal (peak effect at 820 ms) were significantly predicted hEOG tracking. (C) The significant predictors of vEOG tracking were the main effects of speech arousal (peak effect at 1480 ms) and speech valence (peak effect at 100 ms), as well as the interaction effect between valence and mood valence (peak effect at 2020 ms), arousal and mood valence (peak effect at 700 ms), arousal and mood arousal (peak effect at 1540 ms), valence and mood arousal (peak effect at 880 ms).

### 4. Results for emotion models with quadratic term of valence

We initially included both the first (linear) and second (quadratic) degree of the polynomial term for valence ratings in the model to capture potential nonlinear effects. These models revealed several main and interaction effects and were overall better fit than models with only linear valence. However, the inclusion of the quadratic term introduced complex interaction patterns, obscuring a clear interpretation. For clarity and parsimony, we opted to report the model with only linear component of valence in the main text. Here we present the result for quadratic models.

#### 4.1 Predictors of pupil speech tracking

We observed a positive main effect of *speech arousal* in the main model (*β* = 0.04, *p*_FDR_ < .001), suggesting that high arousal speech was tracked stronger than low arousal speech. For *speech valence*, the linear term did not significantly predict pupil-speech tracking, however the quadratic term was significant (*Valence_Linear_*: *β* = −0.004, *p*_uncorrected_ = .22, *p*_FDR_ = .34; *Valence_quadratic_*: *β* = −0.02, p_uncorrected_ < .001, *p*_FDR_ < .001). This suggests that speech tracking is stronger for neutral speech than valenced (positive or negative) speech. In addition, the main effect of *empathy* was significant (*β* = −0.03, *p*_uncorrected_ = .03), however this effect did not remain significant after FDR correction (*p*_FDR_ = .06). No significant main effects were observed for the remaining predictors (see Table S2).

**Table S2.** Summary table for main model predicting pupil-speech MI values. LL = Lower limit, UL = Upper limit, L = Linear term, Q = Quadratic term, A = Arousal dimension, V = Valence dimension. Fixed-effect estimates (β) are reported with confidence intervals (95% CI) computed using the Wald method. *p* values were false discovery rate (FDR) corrected. Significant predictors are highlighted in bold. Blue p-values indicate effects that were also significant in the linear-only model. The model included random intercepts for participants and speech chunks. *\*p* <.05*, **p* <.001

| Predictors | Estimates | 95% CI | | $p_{\text{uncorrected}}$ | $p_{\text{FDR}}$ |
| --- | --- | --- | --- | --- | --- |
|  |  | LL | UL |  |  |
| Intercept | 0.14 | -0.63 | 0.92 | .72 | .79 |
| Arousal | 0.04 | 0.03 | 0.04 | <.001** | <.001** |
| Valence (L) | -0.004 | -0.01 | 0.003 | .22 | .34 |
| Valence (Q) | -0.02 | -0.02 | -0.01 | <.001** | <.001** |
| Mood (A) | 0.01 | -0.02 | 0.04 | .51 | .69 |
| Mood (V) | -0.02 | -0.05 | 0.01 | .25 | .37 |
| Time-in-talk | 0.03 | -0.08 | 0.14 | .61 | .72 |
| Trait empathy | -0.03 | -0.06 | -0.004 | **.03\*** | .06 |
| Pitch | 0.11 | -0.05 | 0.26 | .17 | .29 |
| Loudness | -0.02 | -0.24 | 0.20 | .85 | .85 |
| Talk | -0.09 | -0.61 | 0.44 | .75 | .79 |
| Arousal × Valence (L) | 0.02 | 0.01 | 0.02 | **<.001\*\*** | **<.001\*\*** |
| Arousal × Valence (Q) | -0.01 | -0.01 | -0.004 | **<.001\*\*** | **<.001\*\*** |
| Arousal × Mood (A) | -0.002 | -0.01 | 0.004 | .55 | .69 |
| Arousal × Mood (V) | -0.03 | 0.03 | -0.02 | **<.001\*\*** | **<.001\*\*** |
| Valence (L) × Mood (A) | -0.01 | -0.01 | -0.007 | **<.001\*\*** | **<.001\*\*** |
| Valence (Q) × Mood (A) | 0.03 | 0.02 | 0.03 | **<.001\*\*** | **<.001\*\*** |
| Valence (L) × Mood (V) | -0.01 | -0.01 | -0.000 | **.04\*** | .07 |
| Valence (Q) × Mood (V) | 0.03 | 0.03 | 0.04 | **<.001\*\*** | **<.001\*\*** |

The main model also revealed several interaction effects. First, we found a two-way interaction effect of *speech arousal* × *speech valance*. This suggests that when speech arousal is rated high, positive speech is tracked stronger than negative speech (*speech arousal* × *speech valance*_linear_: *β* = 0.02, *p*_FDR_ < .001), but neutral speech is tracked stronger than both negative and positive speech (*speech arousal* × *speech valance*_quadratic_: *β* = −0.01, *p*_FDR_ < .001). When speech arousal is low, the relationship between valence_linear_ and speech tracking flips, and negative speech is tracked stronger than positive speech.

Second, we found interaction effects between speech emotion (i.e. arousal or valence ratings) and individuals’ own mood (both arousal and valence dimensions). When participants were in a negative mood, they tracked highly arousing speech stronger than low arousing speech, while this relationship was absent for people in a positive mood (*speech arousal × mood valence*: *β* = −0.03, *p*_FDR_ < .001). Additionally, people in a positive mood tracked negative speech more strongly than positive speech (*speech valence_linear_ × mood valence*: *β* = −0.01, *p*_uncorrected_ = .04, *p*_FDR_ = .07) and showed overall stronger tracking of emotional speech than neutral speech (*speech valence_quadratic_ × mood valence*: *β* = 0.03, *p*_FDR_ < .001). In contrast, participants in a negative mood tracked neutral speech stronger than emotional speech.

The arousal dimension of participants’ mood scores only interacted with speech valence (not with speech arousal) to predict pupil speech tracking. Participants who reported high-arousal mood scores showed stronger tracking of negative speech than positive speech (*speech valence_linear_ × mood arousal: β* = −0.01, *p*_FDR_ < .001) and enhanced tracking of emotional speech compared to neutral speech (*speech valence_quadratic_ × mood arousal: β* = 0.03, *p*_FDR_ < .001). Participants in a low-arousal mood tracked neutral speech more strongly than emotional speech.

#### 4.2 Predictors of hEOG speech tracking

The main model with hEOG-speech tracking as outcome revealed that only the main effect of *mood arousal* was significant (*β* = 0.03, *p*_uncorrected_ = .01, *p*_FDR_ = .04), suggesting participants in high arousal mood showed overall stronger tracking of speech. None of the remaining predictors showed a significant main effect (see Table S3).

**Table S3.** Summary table for the mixed effects model predicting hEOG-speech MI value. LL = Lower limit, UL = Upper limit, L = Linear term, Q = Quadratic term A = Arousal dimension V = Valence dimension. Fixed-effect estimates (β) are reported with confidence intervals (95% CI) computed using the Wald method. *p* values were false discovery rate (FDR) corrected. Significant predictors are highlighted in bold. Blue p-values indicate effects that were also significant in the linear-only model. The model included random intercepts for participants and speech chunks. *\*p <.05 **p<.001*

| Predictors | Estimates | 95% CI | | $p_{uncorrected}$ | $p_{FDR}$ |
| --- | --- | --- | --- | --- | --- |
|  |  | LL | UL |  |  |
| Intercept | 0.31 | -0.49 | 1.11 | .45 | .53 |
| Arousal | -0.01 | -0.01 | 0.00 | .09 | .17 |
| Valence (L) | -0.003 | -0.01 | 0.00 | .42 | .53 |
| Valence (Q) | -0.01 | -0.01 | 0.00 | .05 | .12 |
| Mood (A) | 0.03 | 0.01 | 0.06 | .01* | .04* |
| Mood (V) | 0.01 | -0.01 | 0.03 | .43 | .53 |
| Time-in-talk | 0.02 | -0.09 | 0.13 | .73 | .73 |
| Trait empathy | -0.01 | -0.03 | 0.02 | .56 | .62 |
| Pitch | 0.07 | -0.09 | 0.23 | .39 | .53 |
| Loudness | 0.06 | -0.17 | 0.29 | .61 | .64 |
| Talk | -0.22 | -0.76 | 0.33 | .44 | .53 |
| Arousal × Valence (L) | 0.01 | 0.01 | 0.02 | <.001** | <.001** |
| Arousal × Valence (Q) | 0.003 | -0.001 | 0.01 | .11 | .19 |
| Arousal × Mood (A) | 0.02 | 0.02 | 0.03 | <.001** | <.001** |
| Arousal × Mood (V) | -0.02 | -0.03 | -0.01 | <.001** | <.001** |
| Valence (L)× Mood (A) | -0.004 | -0.01 | -0.000 | .08 | .16 |
| Valence (Q)× Mood (A) | -0.01 | -0.01 | 0.004 | <.001** | <.001** |
| Valence (L)× Mood (V) | -0.01 | -0.01 | -0.01 | <.001** | <.001** |
| Valence (Q)× Mood (V) | 0.01 | 0.002 | 0.01 | .002* | .006* |

The interaction between *speech arousal* and *speech valence* was significant (*speech arousal × speech valence_linear_*: *β* = 0.01, *p*_FDR_ < .001). Specifically, hEOG tracking was stronger for negative speech when speech arousal is low, whereas for high-arousal speech, positive speech was tracked more strongly.

The model further revealed significant interaction effects between participants mood and emotional characteristics of speech. Participants in a negative mood tracked highly arousing and positive speech more strongly than low-arousing or negative speech, whereas those in a positive mood exhibited the opposite pattern (*speech arousal × mood valence*: *β* = −0.02, *p*_FDR_ < .001; speech *valence_linear_ × mood valence*: *β* = −0.01, *p*_FDR_ < .001). Additionally, neutral speech was tracked more strongly in a negative mood, while emotional speech was tracked more strongly in a positive mood (speech *valence_quadratic_ × mood valence*: *β* = 0.01, p_uncorrected_ = .002, *p*_FDR_ = 0.006).

When participants were in a low-arousal mood, low-arousal speech was tracked more strongly than high-arousal speech, in contrast, tracking was stronger for high-arousal than for low-arousal speech in a high-arousal mood (*arousal × mood arousal*: *β* = 0.02, *p*_FDR_ < .001). Furthermore, while in a high-arousal mood neutral speech was tracked more strongly than emotionally valenced speech, the opposite was found in a low-arousal mood (*valence_quadratic_ × mood arousal*: *β* = −0.01, *p*_FDR_ < 0.001).

#### 4.3 Predictors of vEOG speech tracking

In the main model predicting vEOG speech tracking, only the main effect of *valence* (*linear*: *β* = −0.01, *p*_uncorrected_ = .05, *p*_FDR_ = .12; *quadratic*: *β* = 0.01, *p*_uncorrected_ = .03, *p*_FDR_ = .07) was significant, suggesting that negative speech was tracked stronger than positive speech (see Table 4).

**Table S4.** Summary table for the mixed effects model predicting vEOG-speech MI values. LL = Lower limit, UL = Upper limit, L = Linear term Q = Quadratic term A = Arousal dimension V = Valence dimension. Fixed-effect estimates (β) are reported with confidence intervals (95% CI) computed using the Wald method. p values were false discovery rate (FDR) corrected. Significant predictors are highlighted in bold. Blue p-values indicate effects that were also significant in the linear-only model. The model included random intercepts for participants and speech chunks. *p <.05 **p<.001.

| Predictors | Estimates | 95% CI | | $p_{\text{uncorrected}}$ | $p_{\text{FDR}}$ |
| --- | --- | --- | --- | --- | --- |
|  |  | LL | UL |  |  |
| Intercept | 0.20 | -0.64 | 1.03 | .65 | .72 |
| Arousal | -0.01 | -0.01 | 0.00 | .05 | .12 |
| Valence (L) | -0.01 | -0.02 | -0.00 | .01* | .04* |
| Valence (Q) | 0.01 | 0.00 | 0.01 | .03* | .07 |
| Mood (A) | 0.02 | -0.01 | 0.05 | .16 | .26 |
| Mood (V) | 0.01 | -0.02 | 0.04 | .46 | .62 |
| Time-in-talk | 0.02 | -0.10 | 0.14 | .77 | .82 |
| Trait empathy | -0.01 | -0.04 | 0.02 | .61 | .72 |
| Pitch | 0.11 | -0.06 | 0.27 | .22 | .32 |
| Loudness | -0.01 | -0.25 | 0.22 | .91 | .91 |
| Talk | -0.14 | -0.70 | 0.43 | .64 | .72 |
| Arousal × Valence (L) | 0.004 | -0.000 | 0.01 | .08 | .14 |
| Arousal × Valence (Q) | -0.01 | -0.01 | -0.002 | .003 | .01* |
| Arousal × Mood (A) | 0.02 | 0.01 | 0.02 | <.001** | <.001** |
| Arousal × Mood (V) | -0.01 | -0.01 | 0.001 | .09 | .15 |
| Valence (L)× Mood (A) | -0.01 | -0.01 | -0.01 | <.001** | <.001** |
| Valence (Q)× Mood (A) | -0.01 | -0.01 | -0.005 | <.001** | <.001** |
| Valence (L)× Mood (V) | 0.02 | 0.01 | 0.02 | <.001** | <.001** |
| Valence (Q)× Mood (V) | -0.004 | -0.01 | 0.000 | .06 | .13 |

We found an interaction effect between speech arousal and speech valence (*arousal × valence_quadratic_*: *β* = −0.01, *p*_FDR_ = 0.01) This interaction effect suggests that emotionally valenced) speech elicited stronger vEOG tracking than neutral speech when the speech was perceived as low in arousal, whereas this pattern diminished for high-arousing speech.

Significant interactions were also observed between participants’ mood and speech emotions. When participants were in a negative mood, tracking of negative speech was stronger than that of positive speech, whereas this pattern reversed for those in a positive mood, who showed stronger tracking of positive than negative speech *(valence_linear_ x mood valence*: *β* = 0.02, *p*_FDR_ < .001).

For mood arousal, participants in a low-arousal mood showed stronger tracking of low-arousal speech compared to high-arousal speech, whereas the opposite pattern was observed for those in a high-arousal mood, who tracked high-arousal speech more strongly (*arousal x mood arousal*: *β* = 0.02, *p*_FDR_ < 0.001). Participants in a low-arousal mood also tracked positive speech more strongly than negative speech (*valence_linear_ x mood arousal β* = −0.01, *p*_FDR_ < 0.001) and showed stronger tracking of valanced speech compared to neutral speech (*valence_quadratic_ x mood arousal: β* = −0.01, *p*_FDR_ < 0.001).

## References

Ahissar, E., & Ahissar, M. (2005). Processing of the temporal envelope of speech. In The auditory cortex: A synthesis of human and animal research (pp. 295–313). Lawrence Erlbaum Associates Publishers.

Arias Sarah, P., Hall, L., Saitovitch, A., Aucouturier, J.-J., Zilbovicius, M., & Johansson, P. (2023). Pupil dilation reflects the dynamic integration of audiovisual emotional speech. Scientific Reports, 13(1), 5507. 10.1038/s41598-023-32133-2

Aston-Jones, G., & Cohen, J. D. (2005). An Integrative Theory of Locus Coeruleus-Norepinephrine Funtion: Adaptive Gain and Optimal Performance. Annual Review of Neuroscience, 28(Volume 28, 2005), 403–450. 10.1146/annurev.neuro.28.061604.135709

Atanasova, T., Gross, J., Rimmele, J., & Keitel, A. (2025). The involvement of endogenous brain rhythms in speech processing. PsyArXiv. 10.31234/osf.io/rukwp_v1

Attaheri, A., Choisdealbha, Á. N., Di Liberto, G. M., Rocha, S., Brusini, P., Mead, N., Olawole-Scott, H., Boutris, P., Gibbon, S., Williams, I., Grey, C., Flanagan, S., & Goswami, U. (2022). Delta- and theta-band cortical tracking and phase-amplitude coupling to sung speech by infants. NeuroImage, 247, 118698. 10.1016/j.neuroimage.2021.118698

Bachorowski, J.-A. (1999). Vocal Expression and Perception of Emotion. Current Directions in Psychological Science, 8(2), 53–57. 10.1111/1467-8721.00013

Baltzell, L. S., Srinivasan, R., & Richards, V. M. (2017). The effect of prior knowledge and intelligibility on the cortical entrainment response to speech. Journal of Neurophysiology, 118(6), 3144–3151. 10.1152/jn.00023.2017

Banse, R., & Scherer, K. R. (1996). Acoustic profiles in vocal emotion expression. Journal of Personality and Social Psychology, 70(3), 614–636. 10.1037//0022-3514.70.3.614

Barr, D. J. (2013). Random effects structure for testing interactions in linear mixed-effects models. Frontiers in Psychology, 4. 10.3389/fpsyg.2013.00328

Bates, D., Mächler, M., Bolker, B., & Walker, S. (2015). Fitting Linear Mixed-Effects Models Using lme4. Journal of Statistical Software, 67, 1–48. 10.18637/jss.v067.i01

Bayer, M., & Schacht, A. (2014). Event-related brain responses to emotional words, pictures, and faces – a cross-domain comparison. Frontiers in Psychology, 5. 10.3389/fpsyg.2014.01106

Belyk, M., & Brown, S. (2014). Perception of affective and linguistic prosody: An ALE meta-analysis of neuroimaging studies. Social Cognitive and Affective Neuroscience, 9(9), 1395–1403. 10.1093/scan/nst124

Benjamini, Y., & Hochberg, Y. (1995). Controlling the False Discovery Rate: A Practical and Powerful Approach to Multiple Testing. Journal of the Royal Statistical Society: Series B (Methodological), 57(1), 289–300. 10.1111/j.2517-6161.1995.tb02031.x

Bestelmeyer, P. E. G., Kotz, S. A., & Belin, P. (2017). Effects of emotional valence and arousal on the voice perception network. Social Cognitive and Affective Neuroscience, 12(8), 1351–1358. 10.1093/scan/nsx059

Bigras, C., Duda, V., & Hébert, S. (2024). Sensory and affective dimensions in loudness perception: Insights from young adults. Hearing Research, 454, 109147. 10.1016/j.heares.2024.109147

Bower, G. H. (1981). Mood and memory. American Psychologist, 36(2), 129–148. 10.1037/0003-066X.36.2.129

Bradley, M. M., Miccoli, L., Escrig, M. A., & Lang, P. J. (2008). The pupil as a measure of emotional arousal and autonomic activation. Psychophysiology, 45(4), 602–607. 10.1111/j.1469-8986.2008.00654.x

Burra, N., Kerzel, D., Munoz Tord, D., Grandjean, D., & Ceravolo, L. (2019). Early spatial attention deployment toward and away from aggressive voices. Social Cognitive and Affective Neuroscience, 14(1), 73–80. 10.1093/scan/nsy100

Ceravolo, L., Frühholz, S., & Grandjean, D. (2016). Modulation of Auditory Spatial Attention by Angry Prosody: An fMRI Auditory Dot-Probe Study. Frontiers in Neuroscience, 10. 10.3389/fnins.2016.00216

Chalas, N., Daube, C., Kluger, D. S., Abbasi, O., Nitsch, R., & Gross, J. (2022). Multivariate analysis of speech envelope tracking reveals coupling beyond auditory cortex. NeuroImage, 258, 119395. 10.1016/j.neuroimage.2022.119395

Chi, T., Ru, P., & Shamma, S. A. (2005). Multiresolution spectrotemporal analysis of complex sounds. The Journal of the Acoustical Society of America, 118(2), 887–906. 10.1121/1.1945807

Clore, G. L., & Huntsinger, J. R. (2007). How emotions inform judgment and regulate thought. Trends in Cognitive Sciences, 11(9), 393–399. 10.1016/j.tics.2007.08.005

Cosme, G., Rosa, P. J., Lima, C. F., Tavares, V., Scott, S., Chen, S., Wilcockson, T. D. W., Crawford, T. J., & Prata, D. (2021). Pupil dilation reflects the authenticity of received nonverbal vocalizations. Scientific Reports, 11(1), 3733. 10.1038/s41598-021-83070-x

Costa, V. D., & Rudebeck, P. H. (2016). More than Meets the Eye: The Relationship between Pupil Size and Locus Coerulus Activity. Neuron, 89(1), 8–10. 10.1016/j.neuron.2015.12.031

Cowie, R., & Cornelius, R. R. (2003). Describing the emotional states that are expressed in speech. Speech Communication, 40(1–2), 5–32. 10.1016/S0167-6393(02)00071-7

de Jong, N. H., & Wempe, T. (2009). Praat script to detect syllable nuclei and measure speech rate automatically. Behavior Research Methods, 41(2), 385–390. 10.3758/BRM.41.2.385

Ding, J., Zhang, X., Liu, J., Hu, Z., Yang, Z., Tang, Y., & Ding, Y. (2025). Entrainment of rhythmic tonal sequences on neural oscillations and the impact on subjective emotion. Scientific Reports, 15(1), 17462. 10.1038/s41598-025-98548-1

Ding, N., Patel, A. D., Chen, L., Butler, H., Luo, C., & Poeppel, D. (2017). Temporal modulations in speech and music. Neuroscience & Biobehavioral Reviews, 81, 181–187. 10.1016/j.neubiorev.2017.02.011

Ding, N., & Simon, J. Z. (2014). Cortical entrainment to continuous speech: Functional roles and interpretations. Frontiers in Human Neuroscience, 8. 10.3389/fnhum.2014.00311

Doelling, K. B., Arnal, L. H., Ghitza, O., & Poeppel, D. (2014). Acoustic landmarks drive delta–theta oscillations to enable speech comprehension by facilitating perceptual parsing. NeuroImage, 85, 761–768. 10.1016/j.neuroimage.2013.06.035

Egidi, G., & Nusbaum, H. C. (2012). Emotional language processing: How mood affects integration processes during discourse comprehension. Brain and Language, 122(3), 199–210. 10.1016/j.bandl.2011.12.008

Eyben, F., Scherer, K. R., Schuller, B. W., Sundberg, J., André, E., Busso, C., Devillers, L. Y., Epps, J., Laukka, P., Narayanan, S. S., & Truong, K. P. (2015). The Geneva Minimalistic Acoustic Parameter Set (GeMAPS) for Voice Research and Affective Computing. IEEE Transactions on Affective Computing, 7(2), 190–202. 10.1109/TAFFC.2015.2457417

Eyben, F., Wöllmer, M., & Schuller, B. (2010). Opensmile: The munich versatile and fast open-source audio feature extractor. Proceedings of the 18th ACM International Conference on Multimedia, 1459–1462. 10.1145/1873951.1874246

Ferré, P., Sánchez-Carmona, A. J., Haro, J., Calvillo-Torres, R., Albert, J., & Hinojosa, J. A. (2025). How does emotional content influence visual word recognition? A meta-analysis of valence effects. Psychonomic Bulletin & Review, 32(2), 570–587. 10.3758/s13423-024-02555-8

Forgas, J. P. (2017). Mood effects on cognition: Affective influences on the content and process of information processing and behavior. In Emotions and affect in human factors and human-computer interaction (pp. 89–122). Academic Press. 10.1016/B978-0-12-801851-4.00003-3

Friston, K. (2010). The free-energy principle: A unified brain theory? Nature Reviews Neuroscience, 11(2), 127–138. 10.1038/nrn2787

Gao, C., Shinkareva, S. V., & Peelen, M. V. (2022). Affective valence of words differentially affects visual and auditory word recognition. Journal of Experimental Psychology: General, 151(9), 2144–2159. 10.1037/xge0001176

Gehmacher, Q., Schubert, J., Schmidt, F., Hartmann, T., Reisinger, P., Rösch, S., Schwarz, K., Popov, T., Chait, M., & Weisz, N. (2024). Eye movements track prioritized auditory features in selective attention to natural speech. Nature Communications, 15(1), 3692. 10.1038/s41467-024-48126-2

Gelman, A., & Hill, J. (2007). Data Analysis Using Regression and Multilevel/Hierarchical Models. Cambridge University Press.

Gerdes, A. B. M., Alpers, G. W., Braun, H., Köhler, S., Nowak, U., & Treiber, L. (2021). Emotional sounds guide visual attention to emotional pictures: An eye-tracking study with audio-visual stimuli. Emotion, 21(4), 679–692. 10.1037/emo0000729

Ghitza, O. (2017). Acoustic-driven delta rhythms as prosodic markers. Language, Cognition and Neuroscience, 32(5), 545–561. 10.1080/23273798.2016.1232419

Gingras, B., Marin, M. M., Puig-Waldmüller, E., & Fitch, W. T. (2015). The Eye is Listening: Music-Induced Arousal and Individual Differences Predict Pupillary Responses. Frontiers in Human Neuroscience, 9. 10.3389/fnhum.2015.00619

Giraud, A.-L., & Poeppel, D. (2012). Cortical oscillations and speech processing: Emerging computational principles and operations. Nature Neuroscience, 15(4), 511–517. 10.1038/nn.3063

Gordon, M. S., & Ancheta, J. (2017). Visual and acoustic information supporting a happily expressed speech-in-noise advantage. The Quarterly Journal of Experimental Psychology, 70(1), 163–178. 10.1080/17470218.2015.1130069

Goudbeek, M., & Scherer, K. (2010). Beyond arousal: Valence and potency/control cues in the vocal expression of emotion. The Journal of the Acoustical Society of America, 128(3), 1322–1336. 10.1121/1.3466853

Grandjean, D., Sander, D., Pourtois, G., Schwartz, S., Seghier, M. L., Scherer, K. R., & Vuilleumier, P. (2005). The voices of wrath: Brain responses to angry prosody in meaningless speech. Nature Neuroscience, 8(2), 145–146. 10.1038/nn1392

Gross, J., Hoogenboom, N., Thut, G., Schyns, P., Panzeri, S., Belin, P., & Garrod, S. (2013). Speech Rhythms and Multiplexed Oscillatory Sensory Coding in the Human Brain. PLOS Biology, 11(12), e1001752. 10.1371/journal.pbio.1001752

Guex, R., Grandjean, D., & Frühholz, S. (2022). Behavioral correlates of temporal attention biases during emotional prosody perception. Scientific Reports, 12(1), 16754. 10.1038/s41598-022-20806-3

Hamann, S. (2012). What can neuroimaging meta-analyses really tell us about the nature of emotion? Behavioral and Brain Sciences, 35(3), 150–152. 10.1017/S0140525X11001701

Hjortkjær, J., Märcher-Rørsted, J., Fuglsang, S. A., & Dau, T. (2020). Cortical oscillations and entrainment in speech processing during working memory load. European Journal of Neuroscience, 51(5), 1279–1289. 10.1111/ejn.13855

Ho, H. T., Schröger, E., & Kotz, S. A. (2015). Selective Attention Modulates Early Human Evoked Potentials during Emotional Face–Voice Processing. Journal of Cognitive Neuroscience, 27(4), 798–818. 10.1162/jocn_a_00734

Hofbauer, L. M., & Rodriguez, F. S. (2023). Emotional valence perception in music and subjective arousal: Experimental validation of stimuli. International Journal of Psychology, 58(5), 465–475. 10.1002/ijop.12922

Hofmann, M. J., Kuchinke, L., Tamm, S., Võ, M. L. H., & Jacobs, A. M. (2009). Affective processing within 1/10th of a second: High arousal is necessary for early facilitative processing of negative but not positive words. Cognitive, Affective, & Behavioral Neuroscience, 9(4), 389–397. 10.3758/9.4.389

Holtze, B., Rosenkranz, M., Bleichner, M., Jaeger, M., & Debener, S. (2023). Eye-Blink Patterns Reflect Attention to Continuous Speech. Advances in Cognitive Psychology, 19(2), 177–200. 10.5709/acp-0387-6

Houston, D., & Haddock, G. (2007). On auditing auditory information: The influence of mood on memory for music. Psychology of Music, 35(2), 201–212. 10.1177/0305735607070303

Ince, R. A. A., Giordano, B. L., Kayser, C., Rousselet, G. A., Gross, J., & Schyns, P. G. (2017). A statistical framework for neuroimaging data analysis based on mutual information estimated via a gaussian copula. Human Brain Mapping, 38(3), 1541–1573. 10.1002/hbm.23471

J Trost, W., Labbé, C., & Grandjean, D. (2017). Rhythmic entrainment as a musical affect induction mechanism. Neuropsychologia, 96, 96–110. 10.1016/j.neuropsychologia.2017.01.004

Jin, P., Zou, J., Zhou, T., & Ding, N. (2018). Eye activity tracks task-relevant structures during speech and auditory sequence perception. Nature Communications, 9(1), 5374. 10.1038/s41467-018-07773-y

Jolliffe, I. T., & Cadima, J. (2016). Principal component analysis: A review and recent developments. Philosophical Transactions. Series A, Mathematical, Physical, and Engineering Sciences, 374(2065), 20150202. 10.1098/rsta.2015.0202

Jürgens, R., Fischer, J., & Schacht, A. (2018). Hot Speech and Exploding Bombs: Autonomic Arousal During Emotion Classification of Prosodic Utterances and Affective Sounds. Frontiers in Psychology, 9, 228. 10.3389/fpsyg.2018.00228

Juslin, P. N., Harmat, L., Barradas, G., Omstedt, G., & Redtzer, V. (2024). Rhythmic entrainment of heart rate as a mechanism for musical emotion induction: A plausible hypothesis in need of evidence? Psychology of Music, 03057356241302809. 10.1177/03057356241302809

Juslin, P. N., & Laukka, P. (2003). Communication of emotions in vocal expression and music performance: Different channels, same code? Psychological Bulletin, 129(5), 770.

Kalashnikova, M., Peter, V., Di Liberto, G. M., Lalor, E. C., & Burnham, D. (2018). Infant-directed speech facilitates seven-month-old infants’ cortical tracking of speech. Scientific Reports, 8(1), 13745. 10.1038/s41598-018-32150-6

Kamiloğlu, R. G., Slocombe, K. E., Haun, D. B. M., & Sauter, D. A. (2020). Human listeners’ perception of behavioural context and core affect dimensions in chimpanzee vocalizations.

Kanske, P., & Kotz, S. A. (2010). Leipzig Affective Norms for German: A reliability study. Behavior Research Methods, 42(4), 987–991. 10.3758/BRM.42.4.987

Kauschke, C., Bahn, D., Vesker, M., & Schwarzer, G. (2019). The Role of Emotional Valence for the Processing of Facial and Verbal Stimuli—Positivity or Negativity Bias? Frontiers in Psychology, 10. 10.3389/fpsyg.2019.01654

Keitel, A., Gross, J., & Kayser, C. (2018). Perceptually relevant speech tracking in auditory and motor cortex reflects distinct linguistic features. PLOS Biology, 16(3), e2004473. 10.1371/journal.pbio.2004473

Keitel, A., Ince, R. A. A., Gross, J., & Kayser, C. (2017). Auditory cortical delta-entrainment interacts with oscillatory power in multiple fronto-parietal networks. NeuroImage, 147, 32–42. 10.1016/j.neuroimage.2016.11.062

Keitel, A., Keitel, C., Alavash, M., Bakardjian, K., Benwell, C. S. Y., Bouton, S., Busch, N. A., Criscuolo, A., Doelling, K. B., Dugue, L., Grabot, L., Gross, J., Hanslmayr, S., Klatt, L.-I., Kluger, D. S., Learmonth, G., London, R. E., Lubinus, C., Martin, A. E.,…Kotz, S. A. (2025). Brain rhythms in cognition—Controversies and future directions (No. arXiv:2507.15639). arXiv. 10.48550/arXiv.2507.15639

Keitel, A., Pelofi, C., Guan, X., Watson, E., Wight, L., Allen, S., Mencke, I., Keitel, C., & Rimmele, J. (2025). Cortical and behavioral tracking of rhythm in music: Effects of pitch predictability, enjoyment, and expertise. Annals of the New York Academy of Sciences, 1546(1), 120–135. 10.1111/nyas.15315

Kim, S. K., & Sumner, M. (2017). Beyond lexical meaning: The effect of emotional prosody on spoken word recognition. The Journal of the Acoustical Society of America, 142(1), EL49–EL55. 10.1121/1.4991328

Kirwan, J., Başkent, D., & Wagner, A. (2025). The Time Course of the Pupillary Response to Auditory Emotions in Pseudospeech, Music, and Vocalizations. Trends in Hearing, 29, 23312165251365824. 10.1177/23312165251365824

Kleiner, M., Denis Pelli, & David Brainard. (2007). What’s new in Psychtoolbox-3?

Koelewijn, T., Shinn-Cunningham, B. G., Zekveld, A. A., & Kramer, S. E. (2014). The pupil response is sensitive to divided attention during speech processing. Hearing Research, 312, 114–120. 10.1016/j.heares.2014.03.010

Koike, K. J., Hurst, M. K., & Wetmore, S. J. (1994). Correlation between the American Academy of Otolaryngology—Head and Neck Surgery Five-Minute Hearing Test and Standard Audiologic Data. Otolaryngology–Head and Neck Surgery, 111(5), 625–632. 10.1177/019459989411100514

Koo, T. K., & Li, M. Y. (2016). A Guideline of Selecting and Reporting Intraclass Correlation Coefficients for Reliability Research. Journal of Chiropractic Medicine, 15(2), 155–163. 10.1016/j.jcm.2016.02.012

Kousta, S.-T., Vinson, D. P., & Vigliocco, G. (2009). Emotion words, regardless of polarity, have a processing advantage over neutral words. Cognition, 112(3), 473–481. 10.1016/j.cognition.2009.06.007

Kuchinke, L., Schneider, D., Kotz, S. A., & Jacobs, A. M. (2011). Spontaneous but not explicit processing of positive sentences impaired in Asperger’s syndrome: Pupillometric evidence. Neuropsychologia, 49(3), 331–338. 10.1016/j.neuropsychologia.2010.12.026

Kuppens, P., Tuerlinckx, F., Russell, J. A., & Barrett, L. F. (2013). The relation between valence and arousal in subjective experience. Psychological Bulletin, 139(4), 917–940. 10.1037/a0030811

Kuznetsova, A., Brockhoff, P. B., & Christensen, R. H. B. (2017). lmerTest Package: Tests in Linear Mixed Effects Models. Journal of Statistical Software, 82, 1–26. 10.18637/jss.v082.i13

Lane, H. (1965). The motor theory of speech perception: A critical review. Psychological Review, 72(4), 275–309. 10.1037/h0021986

Larrouy-Maestri, P., Poeppel, D., & Pell, M. D. (2025). The Sound of Emotional Prosody: Nearly 3 Decades of Research and Future Directions. Perspectives on Psychological Science, 20(4), 623–638. 10.1177/17456916231217722

Laukka, P., Juslin, P., & Bresin, R. (2005). A dimensional approach to vocal expression of emotion. Cognition & Emotion, 19(5), 633–653. 10.1080/02699930441000445

Lehmann, S. J., & Corneil, B. D. (2016). Transient Pupil Dilation after Subsaccadic Microstimulation of Primate Frontal Eye Fields. The Journal of Neuroscience, 36(13), 3765–3776. 10.1523/JNEUROSCI.4264-15.2016

Lesenfants, D., & Francart, T. (2020). The interplay of top-down focal attention and the cortical tracking of speech. Scientific Reports, 10(1), 6922. 10.1038/s41598-020-63587-3

Liu, T., Kotz, S., Criscuolo, A., & Schwartze, M. (2025). The pupil-brain system at rest: Spontaneous pupil fluctuations as markers of neuromodulatory and network dynamic. 10.31234/osf.io/my72t_v1

Lu, L., Ding, Y., Xue, C., & Li, L. (2021). Negative emotions in the target speaker’s voice enhance speech recognition under “cocktail-party” environments. Attention, Perception, & Psychophysics, 83(1), 247–259. 10.3758/s13414-020-02149-1

Lüdecke, D., Bartel, A., Schwemmer, C., Powell, C., Djalovski, A., & Titz, J. (2025). sjPlot: Data Visualization for Statistics in Social Science (Version 2.9.0) [Computer software]. https://cran.r-project.org/web/packages/sjPlot/index.html

Luo, H., & Poeppel, D. (2007). Phase Patterns of Neuronal Responses Reliably Discriminate Speech in Human Auditory Cortex. Neuron, 54(6), 1001–1010. 10.1016/j.neuron.2007.06.004

Madsen, J., & Parra, L. C. (2024). Bidirectional brain-body interactions during natural story listening. Cell Reports, 43(4). 10.1016/j.celrep.2024.114081

Maffei, A., Spironelli, C., & Angrilli, A. (2019). Affective and cortical EEG gamma responses to emotional movies in women with high vs low traits of empathy. Neuropsychologia, 133, 107175. 10.1016/j.neuropsychologia.2019.107175

Majid, A. (2012). Current Emotion Research in the Language Sciences. Emotion Review, 4(4), 432–443. 10.1177/1754073912445827

Martínez-Velázquez, E. S., Ahuatzin González, A. L., Chamorro, Y., & Sequeira, H. (2020). The Influence of Empathy Trait and Gender on Empathic Responses. A Study With Dynamic Emotional Stimulus and Eye Movement Recordings. Frontiers in Psychology, 11, 23. 10.3389/fpsyg.2020.00023

Mathôt, S. (2018). Pupillometry: Psychology, Physiology, and Function. Journal of Cognition, 1(1), 16. 10.5334/joc.18

Mauchand, M., & Zhang, S. (2023). Disentangling emotional signals in the brain: An ALE meta-analysis of vocal affect perception. Cognitive, Affective, & Behavioral Neuroscience, 23(1), 17–29. 10.3758/s13415-022-01030-y

Mayer, J. D., & Gaschke, Y. N. (1988). The experience and meta-experience of mood. Journal of Personality and Social Psychology, 55(1), 102–111. 10.1037//0022-3514.55.1.102

Menn, K. H., Michel, C., Meyer, L., Hoehl, S., & Männel, C. (2022). Natural infant-directed speech facilitates neural tracking of prosody. NeuroImage, 251, 118991. 10.1016/j.neuroimage.2022.118991

Merchie, A., Ranty, Z., Aguillon-Hernandez, N., Aucouturier, J.-J., Wardak, C., & Gomot, M. (2024). Emotional contagion to vocal smile revealed by combined pupil reactivity and motor resonance. Scientific Reports, 14(1), 25043. 10.1038/s41598-024-74848-w

Meyer, L. (2018). The neural oscillations of speech processing and language comprehension: State of the art and emerging mechanisms. European Journal of Neuroscience, 48(7), 2609–2621. 10.1111/ejn.13748

Mullennix, J. W., Bihon, T., Bricklemyer, J., Gaston, J., & Keener, J. M. (2002). Effects of variation in emotional tone of voice on speech perception. Language and Speech, 45(Pt 3), 255–283. 10.1177/00238309020450030301

Myers, B. R., Lense, M. D., & Gordon, R. L. (2019). Pushing the Envelope: Developments in Neural Entrainment to Speech and the Biological Underpinnings of Prosody Perception. Brain Sciences, 9(3), 70. 10.3390/brainsci9030070

Nebi, E., Altmann, T., & Roth, M. (2022). The influence of emotional salience on gaze behavior in low and high trait empathy: An exploratory eye-tracking study. The Journal of Social Psychology, 162(1), 109–127. 10.1080/00224545.2021.2001410

Neves, L., Cordeiro, C., Scott, S. K., Castro, S. L., & Lima, C. F. (2018). High emotional contagion and empathy are associated with enhanced detection of emotional authenticity in laughter. Quarterly Journal of Experimental Psychology (2006), 71(11), 2355–2363. 10.1177/1747021817741800

Nygaard, L. C., & Queen, J. S. (2008). Communicating emotion: Linking affective prosody and word meaning. Journal of Experimental Psychology: Human Perception and Performance, 34(4), 1017–1030. 10.1037/0096-1523.34.4.1017

Obleser, J., & Kayser, C. (2019). Neural Entrainment and Attentional Selection in the Listening Brain. Trends in Cognitive Sciences, 23(11), 913–926. 10.1016/j.tics.2019.08.004

Olano, M. A., Elizalde Acevedo, B., Chambeaud, N., Acuña, A., Marcó, M., Kochen, S., & Alba-Ferrara, L. (2020). Emotional salience enhances intelligibility in adverse acoustic conditions. Neuropsychologia, 147, 107580. 10.1016/j.neuropsychologia.2020.107580

Oliva, M., & Anikin, A. (2018). Pupil dilation reflects the time course of emotion recognition in human vocalizations. Scientific Reports, 8(1), 4871. 10.1038/s41598-018-23265-x

Oostenveld, R., Fries, P., Maris, E., & Schoffelen, J.-M. (2011). FieldTrip: Open Source Software for Advanced Analysis of MEG, EEG, and Invasive Electrophysiological Data. Computational Intelligence and Neuroscience, 2011(1), 156869. 10.1155/2011/156869

Partala, T., & Surakka, V. (2003). Pupil size variation as an indication of affective processing. International Journal of Human-Computer Studies, 59(1), 185–198. 10.1016/S1071-5819(03)00017-X

Patel, S., Scherer, K. R., Björkner, E., & Sundberg, J. (2011). Mapping emotions into acoustic space: The role of voice production. Biological Psychology, 87(1), 93–98. 10.1016/j.biopsycho.2011.02.010

Paulmann, S. (2016). Chapter 88—The Neurocognition of Prosody. In G. Hickok & S. L. Small (Eds), Neurobiology of Language (pp. 1109–1120). Academic Press. 10.1016/B978-0-12-407794-2.00088-2

Paulmann, S., Ott, D. V. M., & Kotz, S. A. (2011). Emotional Speech Perception Unfolding in Time: The Role of the Basal Ganglia. PLOS ONE, 6(3), e17694. 10.1371/journal.pone.0017694

Paulmann, S., Titone, D., & Pell, M. D. (2012). How emotional prosody guides your way: Evidence from eye movements. Speech Communication, 54(1), 92–107. 10.1016/j.specom.2011.07.004

Peelle, J. E., Gross, J., & Davis, M. H. (2013). Phase-Locked Responses to Speech in Human Auditory Cortex are Enhanced During Comprehension. Cerebral Cortex, 23(6), 1378–1387. 10.1093/cercor/bhs118

Pell, M. D., & Kotz, S. A. (2021). Comment: The Next Frontier: Prosody Research Gets Interpersonal. Emotion Review, 13(1), 51–56. 10.1177/1754073920954288

Pessoa, L. (2008). On the relationship between emotion and cognition. Nature Reviews Neuroscience, 9(2), 148–158. 10.1038/nrn2317

Pfeffer, T., Keitel, C., Kluger, D. S., Keitel, A., Russmann, A., Thut, G., Donner, T. H., & Gross, J. (2022). Coupling of pupil- and neuronal population dynamics reveals diverse influences of arousal on cortical processing. eLife, 11, e71890. 10.7554/eLife.71890

Pinheiro, A. P., Barros, C., Dias, M., & Niznikiewicz, M. (2017). Does emotion change auditory prediction and deviance detection? Biological Psychology, 127, 123–133. 10.1016/j.biopsycho.2017.05.007

R Core Team. (2024). [Computer software].

Rigoulot, S., & Pell, M. D. (2012). Seeing Emotion with Your Ears: Emotional Prosody Implicitly Guides Visual Attention to Faces. PLOS ONE, 7(1), e30740. 10.1371/journal.pone.0030740

Rimmele, J. M., Poeppel, D., & Ghitza, O. (2021). Acoustically Driven Cortical δ Oscillations Underpin Prosodic Chunking. Eneuro, 8(4), ENEURO.0562-20.2021. 10.1523/ENEURO.0562-20.2021

Rimmele, J. M., Zion Golumbic, E., Schröger, E., & Poeppel, D. (2015). The effects of selective attention and speech acoustics on neural speech-tracking in a multi-talker scene. Cortex: A Journal Devoted to the Study of the Nervous System and Behavior, 68, 144–154. 10.1016/j.cortex.2014.12.014

Rosen, S. (1992). Temporal information in speech: Acoustic, auditory and linguistic aspects (p. 79). Clarendon Press/Oxford University Press.

Russell, J. A. (1980). A circumplex model of affect. Journal of Personality and Social Psychology, 39(6), 1161–1178. 10.1037/h0077714

Saint-Georges, C., Chetouani, M., Cassel, R., Apicella, F., Mahdhaoui, A., Muratori, F., Laznik, M.-C., & Cohen, D. (2013). Motherese in Interaction: At the Cross-Road of Emotion and Cognition? (A Systematic Review). PLOS ONE, 8(10), e78103. 10.1371/journal.pone.0078103

Schirmer, A., & Kotz, S. A. (2006). Beyond the right hemisphere: Brain mechanisms mediating vocal emotional processing. Trends in Cognitive Sciences, 10(1), 24–30. 10.1016/j.tics.2005.11.009

Schirmer, A., Kotz, S. A., & Friederici, A. D. (2005). On the role of attention for the processing of emotions in speech: Sex differences revisited. Cognitive Brain Research, 24(3), 442–452. 10.1016/j.cogbrainres.2005.02.022

Schröder, M. (2001). Emotional speech synthesis: A review. 7th European Conference on Speech Communication and Technology (Eurospeech 2001), 561–564. 10.21437/Eurospeech.2001-150

Schubert, J., Gehmacher, Q., Schmidt, F., Hartmann, T., & Weisz, N. (2025). Prediction tendency, eye movements, and attention in a unified framework of neural speech tracking (p. 2023.06.27.546746). bioRxiv. 10.1101/2023.06.27.546746

Scott, S. K., McGettigan, C., & Eisner, F. (2009). A little more conversation, a little less action—Candidate roles for the motor cortex in speech perception. Nature Reviews Neuroscience, 10(4), 295–302. 10.1038/nrn2603

Smith, Z. M., Delgutte, B., & Oxenham, A. J. (2002). Chimaeric sounds reveal dichotomies in auditory perception. Nature, 416(6876), 87–90. 10.1038/416087a

Song, J., & Iverson, P. (2018). Listening effort during speech perception enhances auditory and lexical processing for non-native listeners and accents. Cognition, 179, 163–170. 10.1016/j.cognition.2018.06.001

Spreng, R. N., McKinnon, M. C., Mar, R. A., & Levine, B. (2009). The Toronto Empathy Questionnaire: Scale Development and Initial Validation of a Factor-Analytic Solution to Multiple Empathy Measures. Journal of Personality Assessment, 91(1), 62–71. 10.1080/00223890802484381

Teoh, E. S., Cappelloni, M. S., & Lalor, E. C. (2019). Prosodic pitch processing is represented in delta-band EEG and is dissociable from the cortical tracking of other acoustic and phonetic features. European Journal of Neuroscience, 50(11), 3831–3842. 10.1111/ejn.14510

Themistocleous, C. (2025). Linguistic and emotional prosody: A systematic review and ALE meta-analysis. Neuroscience & Biobehavioral Reviews, 175, 106210. 10.1016/j.neubiorev.2025.106210

Trainor, L. J., Austin, C. M., & Desjardins, R. N. (2000). Is Infant-Directed Speech Prosody a Result of the Vocal Expression of Emotion? Psychological Science, 11(3), 188–195. 10.1111/1467-9280.00240

Urai, A. E., Braun, A., & Donner, T. H. (2017). Pupil-linked arousal is driven by decision uncertainty and alters serial choice bias. Nature Communications, 8(1), 14637. 10.1038/ncomms14637

Van Berkum, J. J. A., de Goede, D., van Alphen, P., Mulder, E., & Kerstholt, J. H. (2013). How robust is the language architecture? The case of mood. Frontiers in Psychology, 4. 10.3389/fpsyg.2013.00505

Vuilleumier, P. (2005). How brains beware: Neural mechanisms of emotional attention. Trends in Cognitive Sciences, 9(12), 585–594. 10.1016/j.tics.2005.10.011

Wallmark, Z., Deblieck, C., & Iacoboni, M. (2018). Neurophysiological Effects of Trait Empathy in Music Listening. Frontiers in Behavioral Neuroscience, 12. 10.3389/fnbeh.2018.00066

Wang, C.-A., & Munoz, D. P. (2015). A circuit for pupil orienting responses: Implications for cognitive modulation of pupil size. Current Opinion in Neurobiology, 33, 134–140. 10.1016/j.conb.2015.03.018

Weninger, F., Eyben, F., Schuller, B. W., Mortillaro, M., & Scherer, K. R. (2013). On the Acoustics of Emotion in Audio: What Speech, Music, and Sound have in Common. Frontiers in Psychology, 4. 10.3389/fpsyg.2013.00292

Wurm, L. H., Vakoch, D. A., Strasser, M. R., Calin-Jageman, R., & Ross, S. E. (2001). Speech perception and vocal expression of emotion. Cognition and Emotion, 15(6), 831–852. 10.1080/02699930143000086

Yang, W., Makita, K., Nakao, T., Kanayama, N., Machizawa, M. G., Sasaoka, T., Sugata, A., Kobayashi, R., Hiramoto, R., Yamawaki, S., Iwanaga, M., & Miyatani, M. (2018). Affective auditory stimulus database: An expanded version of the International Affective Digitized Sounds (IADS-E). Behavior Research Methods, 50(4), 1415–1429. 10.3758/s13428-018-1027-6

Zekveld, A. A., Koelewijn, T., & Kramer, S. E. (2018). The Pupil Dilation Response to Auditory Stimuli: Current State of Knowledge. Trends in Hearing, 22, 2331216518777174. 10.1177/2331216518777174

Zekveld, A. A., Kramer, S. E., & Festen, J. M. (2010). Pupil Response as an Indication of Effortful Listening: The Influence of Sentence Intelligibility. Ear and Hearing, 31(4), 480. 10.1097/AUD.0b013e3181d4f251

